# A multilayered post-GWAS assessment on genetic susceptibility to pancreatic cancer

**DOI:** 10.1101/2020.02.11.941351

**Authors:** E López de Maturana, JA Rodríguez, L Alonso, O Lao, E Molina-Montes, I Martín-Antoniano, P Gómez-Rubio, RT Lawlor, A Carrato, M Hidalgo, M Iglesias, X Molero, M Löhr, CW Michalski, J Perea, M O’Rorke, VM Barberà, A Tardón, A Farré, L Muñoz-Bellvís, T Crnogorac-Jurcevic, E Domínguez-Muñoz, T Gress, W Greenhalf, L Sharp, L Arnes, Ll Cecchini, J Balsells, E Costello, L Ilzarbe, J Kleeff, B Kong, M Márquez, J Mora, D O’Driscoll, A Scarpa, W Ye, J Yu, PanGenEU Investigators, M García-Closas, M Kogevinas, N Rothman, D Silverman, SBC/EPICURO Investigators, D Albanes, AA Arslan, L Beane-Freeman, PM Bracci, P Brennan, B Bueno-de-Mesquita, J Buring, F Canzian, M Du, S Gallinger, JM Gaziano, PJ Goodman, M Gunter, L LeMarchand, D Li, RE Neale, U Peters, GM Petersen, HA Risch, MJ Sánchez, XO Shu, MD Thornquist, K Visvanathan, W Zheng, S Chanock, D Easton, BM Wolpin, RZ Stolzenberg-Solomon, AP Klein, LT Amundadottir, MA Marti-Renom, FX Real, N Malats

**Author notes:** Equal contributions (E López de Maturana, JA Rodríguez). **Correspondence to:** Núria Malats, Genetic and Molecular Epidemiology Group, Spanish National Cancer Research Center (CNIO), C/Melchor Fernandez Almagro 3, 28029-Madrid, Spain; and Marc A. Martí-Renom, Structural Genomics Group, Centre Nacional d’Anàlisi Genòmica - Centre de Regulació Genòmica (CNAG-CRG), Baldiri Reixac 4, 08028-Barcelona.

## Abstract

Pancreatic cancer (PC) is a complex disease in which both non-genetic and genetic factors interplay. To-date, 40 GWAS hits have been associated with PC risk in individuals of European descent, explaining 4.1% of the phenotypic variance. Here, we complemented a classical new PC GWAS (1D) with spatial autocorrelation analysis (2D) and Hi-C maps (3D) to gain additional insight into the inherited basis of PC. *In-silico* functional analysis of public genomic information allowed prioritization of potentially relevant candidate variants. We replicated 17/40 previous PC-GWAS hits and identified novel variants with potential biological functions. The spatial autocorrelation approach prioritized low MAF variants not detected by GWAS. These were further expanded via 3D interactions to 54 target regions with high functional relevance. This multi-step strategy, combined with an in-depth *in-*silico functional analysis, offers a comprehensive approach to advance the study of PC genetic susceptibility and could be applied to other diseases.

## INTRODUCTION

Pancreatic cancer (PC) has a relatively low incidence but it is one of the deadliest tumors. In Western countries, PC ranks fourth among cancer-related deaths with 5-year survival of 3% in Europe^1-3^. In the last decades, progress in the management of patients with PC has been meagre. In addition, mortality is rising^2^ and it is estimated that PC will become the second cause of cancer-related deaths in the United States by 2030^4^.

PC is a complex disease in which both genetic and non-genetic factors participate. However, relatively little is known about its etiologic and genetic susceptibility background. In comparison with other major cancers, fewer genome-wide association studies (GWAS) have been carried out and the number of patients included in them is relatively small (N=9,040). According to the GWAS Catalog, (January 2019)^5^, 40 common germline variants associated with PC risk have been identified in 32 loci in individuals of European descent^6-11^. However, these variants only explain 4.1% of the phenotypic variance for PC^12^. More importantly, given the challenges in performing new PC case-control studies with adequate clinical, epidemiological, and genetic information, the field is far from reaching the statistical power that has been achieved in other more common cancers such as breast, colorectal, or prostate cancers with >100,000 subjects included in GWAS, yielding a much larger number of genetic variants associated with them^5^.

Current GWAS methodology relies on setting a strict statistical threshold of significance (*p*-value=5×10^−8^) and on replication in independent studies. This approach has been successful in minimizing false positive hits at the expense of discarding variants that may be truly associated with the disease (false negatives) displaying association *p*-values not reaching genome-wide significance after multiple testing correction or not being replicated in independent populations. The “simple” solution to this problem is to increase the number of subjects. However, it will take considerable time for PC GWAS studies to reach the sample size achieved in other tumors and the funding climate for replication studies is extremely weak. While a meta-analysis based on available datasets provides an alternative strategy for novel variant identification, this approach may introduce heterogeneity because studies differ regarding methods, data quality, testing strategies, genetic background of the included individuals (e.g., population substructure), and study design, factors that can lead to lack of replicability. Therefore, we are faced with the need of exploring alternative approaches to substantiate findings of putative genetic risk variants not fulfilling conventional GWAS criteria.

Here, we build upon one of the largest epidemiological PC case-control studies with extensive standardized clinical and epidemiological annotation and expand the findings of a classical GWAS to include novel strategies for risk-variant discovery. First, we used the Local Moran’s Index (LMI)^13^, an approach that is widely applied in geospatial statistics. In its original application to geographic two-dimensional analysis, LMI identifies the existence of relevant clusters in the spatial arrangement of a variable, highlighting points closely surrounded by others with similar values, allowing the identification of “hot spots”. In our genomic application, we computed local indexes of spatial (genomic) autocorrelation to identify clusters of SNPs based on their similar magnitudes of association (odds ratio, OR) weighted by their genomic distance as measured by linkage disequilibrium (LD). By capturing LD structures of nearby SNPs, LMI leverages the values of SNPs with low minor allele frequencies (MAF) that conventional GWAS fail to assess properly. In this regard, LMI offers a novel opportunity to identify potentially relevant new set of genomic candidates associated with PC genetic susceptibility.

In addition, we have taken advantage of recent advances in 3D genomic analyses providing insights into the spatial relationship of regulatory elements and their target genes. Since GWAS have largely identified variants present in non-coding regions of the genome, a challenge has been to ascribe such variants to the corresponding regulated genes, which may lie far away in the genomic sequence. Chromosome Conformation Capture experiments (3C-related techniques)^14^ can provide insight into the biology and function underlying previously “unexplained” hits^15,16^.

High-throughput technologies have produced large amounts of publicly-available data from cell types and tissues. Given the hypothesis-free nature of GWAS, the aforementioned resources represent a valuable approach to validate prioritized variants using novel criteria, as well as for functional interpretation of genetic findings.

The combined use of conventional GWAS (1D) analysis with LMI (2D) and 3D genomic approaches has allowed enhancing the discovery of novel candidate variants involved in PC (**Figure 1**). Importantly, several of the new variants are located in genes relevant to the biology and function of pancreatic epithelial cells.

**Figure 1.**
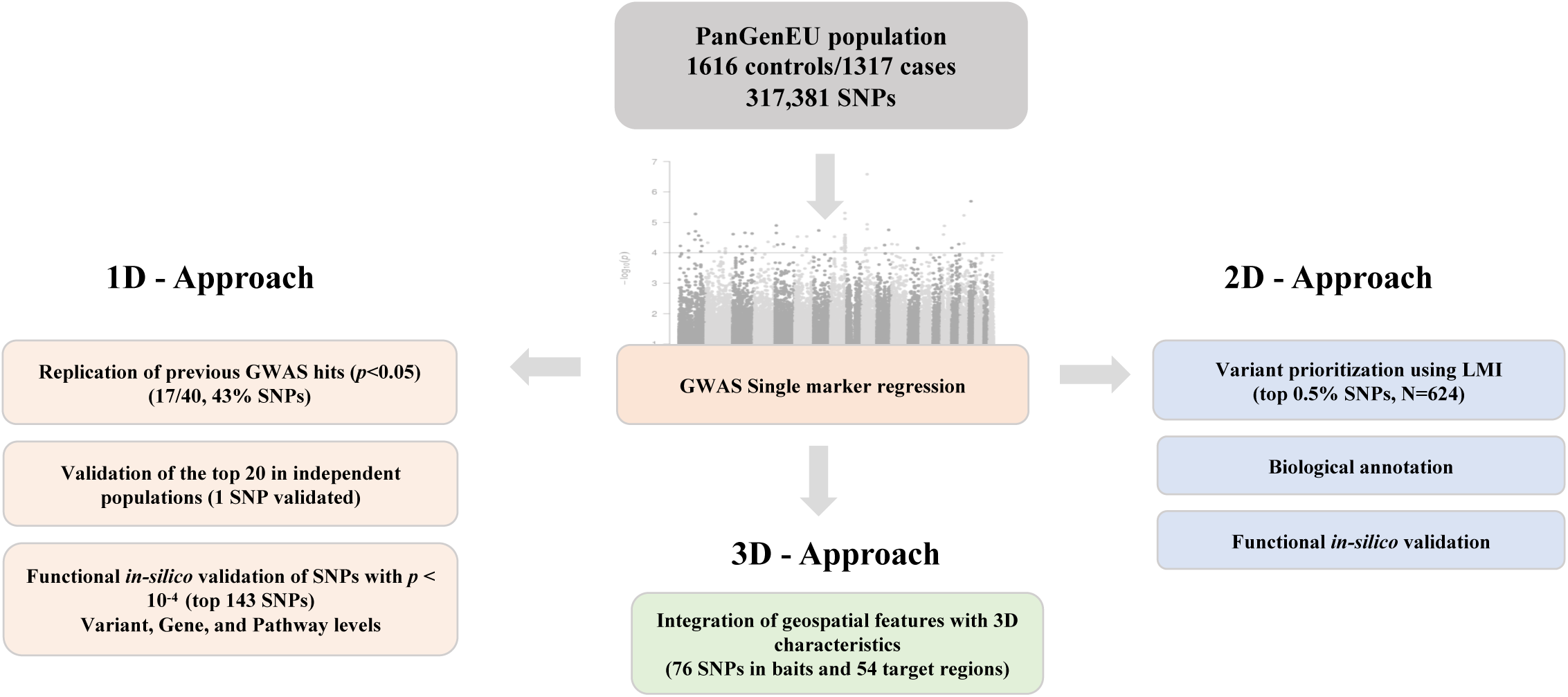
Overview of the approaches adopted in this study to identify new pancreatic cancer susceptibility regions.

## RESULTS

### 1D Approach: PanGenEU GWAS - Single marker association analyses

We performed a GWAS including data from 1,317 patients diagnosed with PC (cases) and 1,616 control individuals from European countries. In addition to all genotyped SNPs that passed the QC procedure, we included imputed data for the previously reported PC-associated hits not genotyped in OncoArray-500K; the 1000G Phase3 (Haplotype release date October 2014) being used as reference^17^. In all, 317,270 SNPs were tested (**Figure S1)** with little evidence of genomic inflation (**Figure S2**).

#### Replication of previously reported GWAS hits

Of the 40 previously GWAS-discovered variants associated with PC risk in European ancestry populations^5^, 17 (42.5%) were replicated with nominal *p-*values<0.05. For all 17, the associations were in the same direction as in the primary reports (**Table S1**). Among them, we replicated *NR5A2*-rs2816938 and *NR5A2*-rs3790844. Furthermore, we observed significant associations for seven additional variants tagging *NR5A2* previously reported in the literature^7-10,18^. At the GWAS significance level, we also replicated the GWAS hits *LINC00673*-rs7214041^11^ and *TERT*-rs2736098^8,11^.

#### Validation of the top 20 PanGenEU GWAS hits in independent populations

The risk estimates of the top 20 variants in the PanGenEU GWAS were included in the meta-analyis with those derived from PanScanI+II, PanScan III, and PanC4 consortia GWAS, representing a total of 10,357 cases and 14,112 controls (**Table S2**). PanGenEU GWAS identified a new variant in *NR5A2* associated with PC (*NR5A2*-rs3790840, metaOR=1.23, *p-*value=5.91×10^−6^) which is in moderate LD with *NR5A2*-rs4465241 (*r*^*2*^=0.45, metaOR=0.81, *p-*value=3.27×10^−10^) and had previously been reported in a GWAS pathway analysis^18^. *NR5A2*-rs3790840 remained significant (*p-*value<0.05) when conditioned on *NR5A2-*rs4465241, on *NR5A2-*rs3790844 plus *NR5A2-*rs2816938, and even on the 13 *NR5A2* GWAS hits reported in the literature, indicating that *NR5A2*-rs3790840 is a new, distinct, PC risk signal. Using SKAT-O (seqMeta R package), we performed a gene-based association analysis considering all significant *NR5A2* hits plus *NR5A2*-rs3790840; the *NR5A2-*based association results were significant (*p-* value=8.9×10^−4^). Furthermore, in a case-only analysis conducted within the PanGenEU study, *NR5A2* variation was also associated with diabetes (*p-*value=6.0×10^−3^), suggesting an interaction between both factors in relation to PC risk.

### Post-GWAS Functional in-silico analyses

#### Assessment of potential functionality of the variants

We expanded the primary assessment by performing a systematic *in silico* functional analysis of SNPs with GWAS *p*-values<1×10^−4^ (N=143) at the variant, gene, and pathway levels (**Figure S3**). The potential functionality of the most relevant SNPs, according to the features considered at all levels, is summarized in **Supplementary Material and Table S3**.

Among the functionally suggestive variants, we highlight those in *CASC8* (8q24.21) (**Figure 2**): 27 variants with *p*-values <1×10^−4^ organized in four LD-blocks were identified. The largest block contained 11 variants (*r*^*2*^=0.87-1). For 8 of them, the ORs of the association alleles were below unity. *CASC8* codes for a non-protein coding RNA overexpressed in tumor vs normal pancreatic tissue (Log2FC=1.25, *p-* value=2.29×10^−56^). All *CASC8* variants were associated with differential leukocyte methylation (mQTL) of *RP11-382A18.1-*cg25220992 in our PanGenEU population sample. Moreover, 20 of them were also associated with differential methylation of cg03314633, also in *RP11-382A18.1*. Twenty-three of the variants overlapped with at least one histone mark in either endocrine or exocrine pancreatic tissue. Two of these hits have been previously associated with other cancers: *CASC8-*rs1562430 (breast, colorectal, and stomach) and *CASC8-*rs2392780 (breast). None of the *CASC8* hits were in LD with *CASC11*-rs180204, a GWAS hit previously associated with PC risk, which is ∼205 Kb downstream^10^. *CASC8* also overlaps with a PC-associated lncRNA^19^, suggesting that genetic variants in *CASC8* may contribute to the transcriptional program of pancreatic tumor cells. Moreover, 5% of PC tumors catalogued in cBioPortal had alterations in *CASC8* (37 cases showed gene amplifications and one sample presented a fusion). Alterations in *CASC8* significantly co-occur with alterations in *TG* (adjusted *p*-values*<*0.001), also associated with PC in our GWAS, which is located downstream.

**Figure 2.**
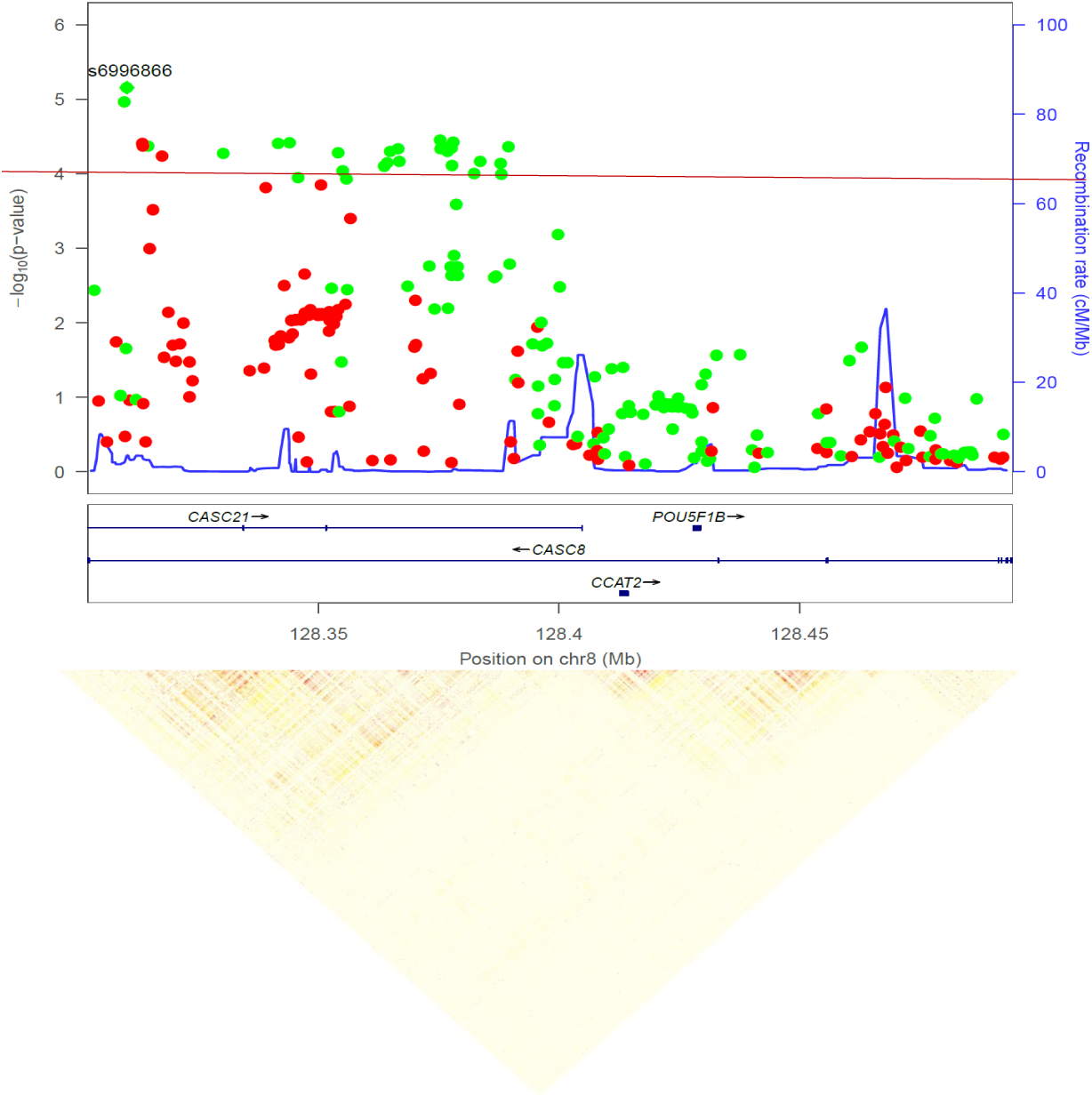
Zoom plot of the 8q24.21 CASC8 (cancer Susceptibility 8) region and linkage disequilibrium pattern of the PanGenEU GWAS prioritized variants.

Three of the variants prioritized for *in-silico* analysis are located in genes involved in pancreatic function: rs1220684 is in *SEC63*, coding for a protein involved in endoplasmic reticulum function and ER stress response^20^; rs7212943, a putative regulatory variant, is in *NOC2/RPH3AL*, a gene involved in exocytosis in exocrine and endocrine cells^21^; and rs4383344 is in *SCTR*, which encodes for the secretin receptor, selectively expressed in the exocrine pancreas and involved in production and indirectly in regulation of bicarbonate, electrolyte, and volume secretion in ductal cells. Interestingly, secretin regulation is affected by *H. pylori* which has been suggested a PC risk factor^22^. High expression of *SCTR* has also been reported in PC^23^.

#### Gene set enrichment analyses

When considering the 81 genes harboring the 143 SNPs prioritized as described above, 6 chromosomal regions were significantly enriched (**Table S4**). Moreover, a gene-set enrichment analysis was performed for the gene-trait associations reported in the GWAS Catalog resulting in 29 traits (**Table S4**). The most relevant GWAS traits with significant enrichment were ‘Pancreatic cancer’, ‘Uric acid levels’, ‘Major depressive disorder’ and ‘Obesity-related traits’, in addition to ‘Lung adenocarcinoma’, ‘Lung cancer’, and ‘Prostate cancer’ traits. We also performed a network analysis using the *igraph* R package^24^ to visualize the relationships between the enriched GWAS traits and the prioritized genes. Twelve densely connected subgraphs were identified via random walks (**Figure 3**). Interestingly, ‘pancreatic cancer’ and ‘uric acid levels’ GWAS traits were connected through *NR5A2*, which is also linked to ‘chronic inflammatory diseases’ and ‘lung carcinoma’ traits. *NR5A2* is an important regulator of pancreatic differentiation and inflammation in the pancreas^25^.

**Figure 3.**
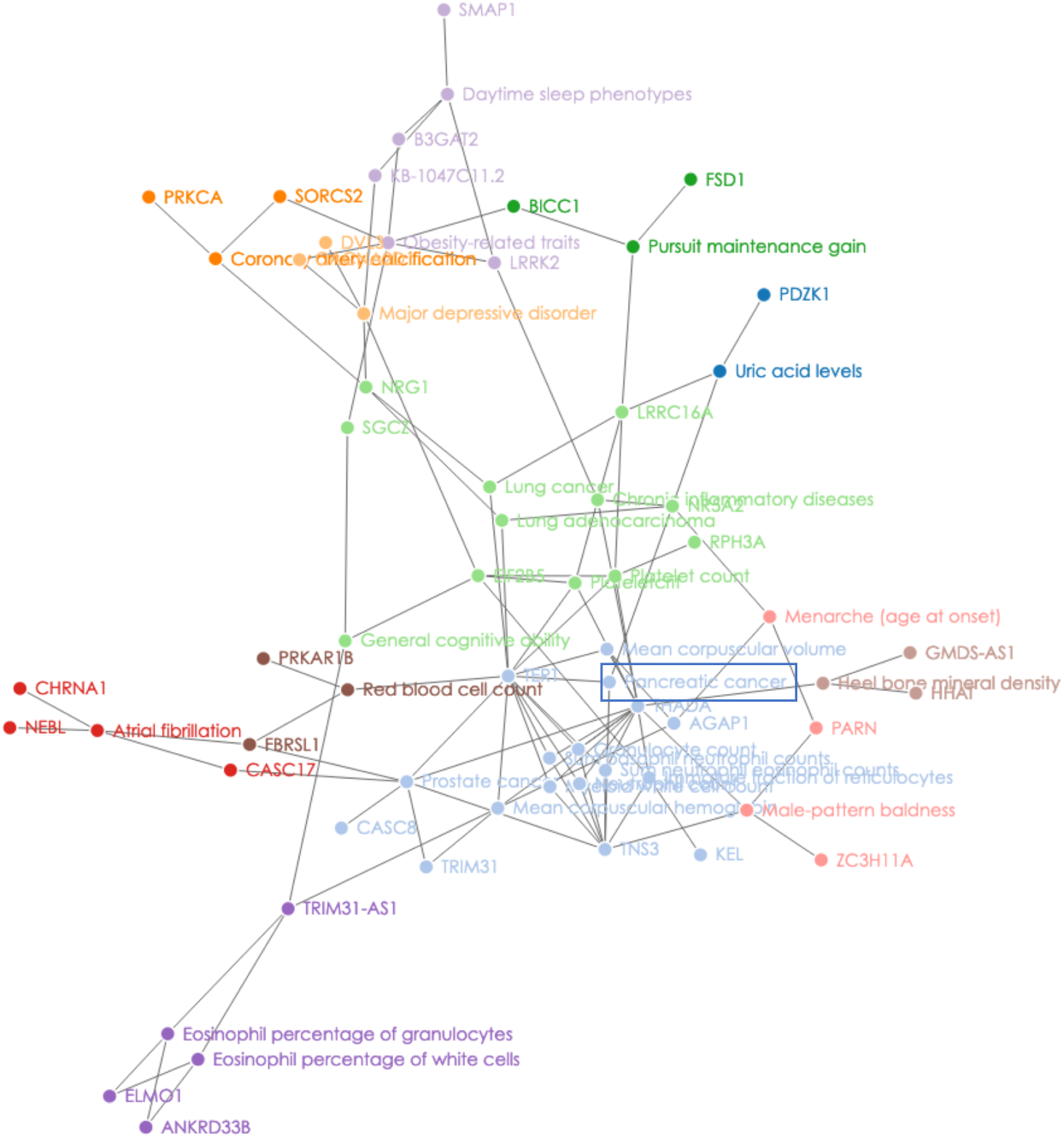
Network of traits in GWAS catalog enriched with the genes prioritized in the PanGenEU GWAS.

#### Pathway enrichment analyses

A total of 112 Gene Ontology (GO) terms according to their biological function (GO:BP) (adjusted *p*-values<0.05, with minimum of three genes overlapping), seven GO terms according to their cellular components (GO:CC) and 11 terms according to their molecular functions (GO:MF) were significantly enriched with the prioritized genes (**Table S4**). Interestingly, GO terms relevant to exocrine pancreatic function were overrepresented. Three KEGG pathways were significantly enriched with ≥2 genes from our prioritized set (**Table S4**); among them are: “Glycosaminoglycan biosynthesis heparan sulfate” (*adj*-*p*=3.86×10^−3^), “ERBB signaling pathway” (*adj*-*p*=3.73×10^−2^) and “Melanogenesis” (*adj*-*p*=3.73×10^−2^). Interconnections between the three significant KEGG pathways after gene enrichment were explored using the *Pathway*-*connector* webtool (**Figure S4**), which also found six complementary pathways: ‘Tyrosine metabolism’, ‘Metabolic pathways’, ‘Glycolysis/Gluconeogenesis’, ‘Glycerolipid metabolism’, ‘PI3K-Akt signaling pathway’, ‘mTOR signaling pathway’.

### 2D-Approach: Integration of geospatial features

#### Variant prioritization using LMI

We scaled up from the single-SNP (1D) to the genomic region (2D) association analysis by considering both genomic distance (LD) between variants and association magnitude (OR). We calculated a LMI score (see Methods) for 98.8% of the SNPs in our dataset, as 1.2% of the SNPs were not genotyped in the 1000 G (Phase 3, v1) reference data set^17^ or had a MAF<1% in the CEU European population (n=85 individuals, phase 1, version 3). We selected those SNPs with positive LMI or within the top 50% of OR values. This filter resulted in a final set of 102,146 SNPs. The LMI scores and *p*-values for these variants showed a direct correlation (Spearman *r*=0.62; *p-*value*=*2.2×10^−16^, **Figure 4**). Next, an LMI-enriched variant set was generated by selecting the top 0.5% of SNPs according to their LMI scores, which included 29 out of the 143 SNPs selected through their GWAS *p-*values. Finally, a combined SNP set was generated by adding the remaining 114 SNPs prioritized in the 1D approach to the LMI-enriched dataset, resulting in 624 SNPs (**Figure 4**). To assess the versatility of LMI, we ran two benchmarks on the MAFs and the ORs, both confirming the potential to prioritize SNPs (**Supplementary Material**). We compared the MAF distribution between the GWAS-prioritized and the LMI-selected SNPs. Notably, LMI-SNPs were mainly variants with low MAF (<0.1). (**Figure S5**). After excluding correlated SNPs among the 143 GWAS-SNPs and the 624 LMI-SNPs by LD (*r*^*2*^ < 0.2 to consider independent loci; Methods), we obtained 97 and 248 independent signals, respectively. Average MAF for the GWAS-prioritized variants was 0.24 (SD=0.13), compared with 0.07 (SD=0.03) for the top-rank LMI-SNPs. This result emphasizes that statistical significance for GWAS-SNPs is largely dependent on MAF and the statistical power of the study, highlighting this as a major limitation of classical GWAS analyses. LMI captured a new dimension of signals independent from MAF (**Figure S5**). In line with the above observation, the average OR for the LMI-SNPs was significantly higher than that for the GWAS-SNPs (1.46 *vs*. 1.32, respectively, Wilcoxon statistic *p-* value*=*1.63×10^−10^). Altogether, these results support the notion that LMI is more sensitive to detect candidate SNPs with lower MAFs but relevant effect sizes.

**Figure 4.**
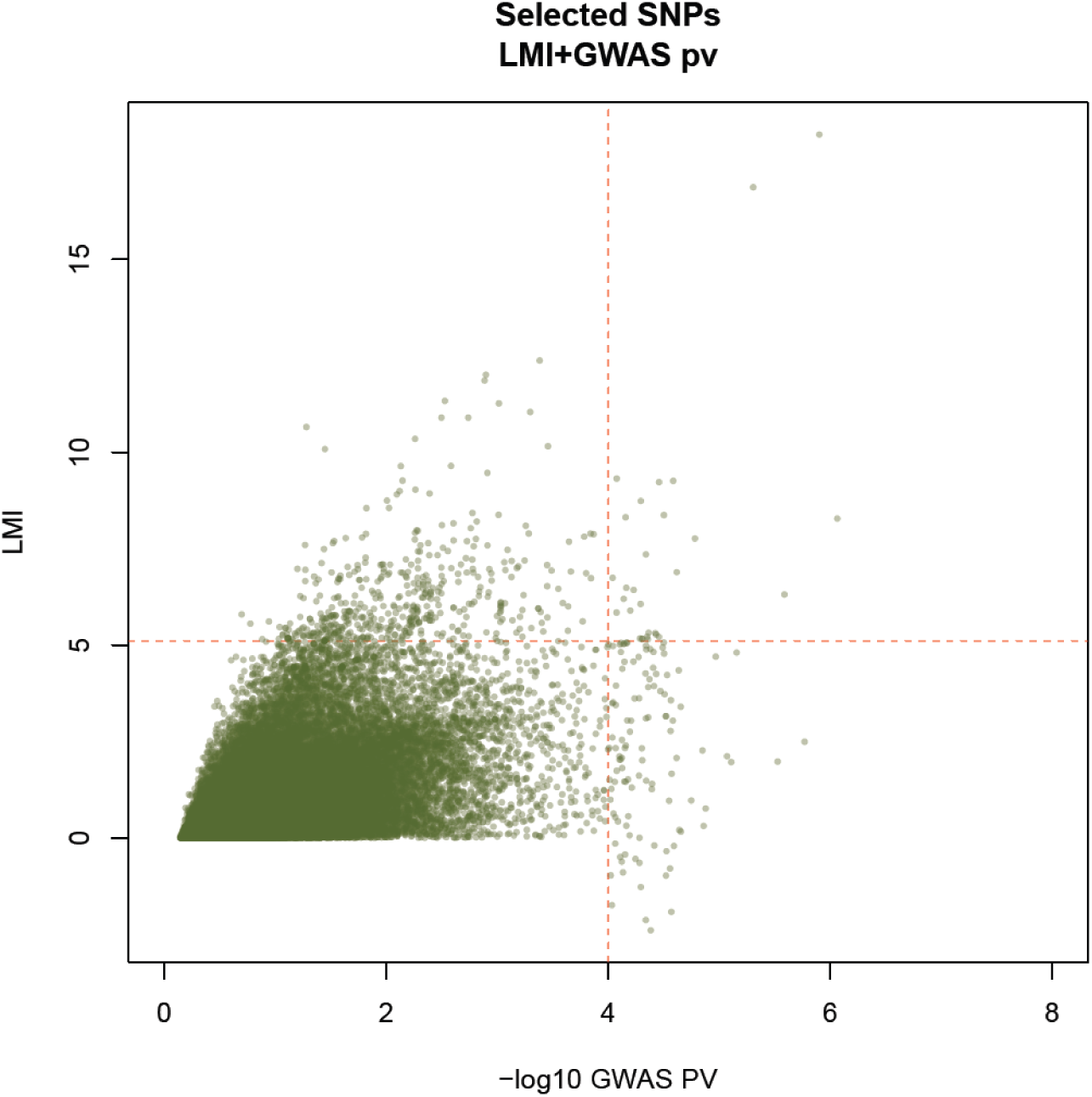
Scatterplot of the local Moran’s index (LMI) obtained in the 2D approach and the –log10 *p*-value obtained in the GWAS analysis (1D approach).

#### Biological annotation of LMI-regions

Variants prioritized according to both LMI and GWAS *p-value* (N=624) were annotated to 338 genes using annotatePeaks.pl HOMER script^26^ (**Table S5**). The two top LMI-SNPs were also captured by the GWAS approach. They map, respectively, to intronic sequences of *MINDY1* (*p-value*= 1.26×10^−6^) and *SETDB1* (*p-value*=4.94×10^−6^). Importantly, among the top SNPs identified by LMI was *BCAR1-*rs7190458, a variant with a relevant role in PC^27^ reported in two previous GWAS^8,11^. An additional SNP (rs13337397) in the first exon of *BCAR1*, and in low LD with *BCAR1-*rs7190458 (*r*^*2*^=0.36), was also prioritized by LMI. This SNP is intergenic to *CTRB1-2* and *BCAR1* (**Figure 5**). While *BCAR1* is ubiquitous, *CTRB1-2* is expressed exclusively in the exocrine pancreas and genetic variation therein has been previously associated with alcoholic pancreatitis^28^ and type-2 diabetes^29,30^. The expression of both genes is reduced in tumors vs. normal tissue^31^.

**Figure 5.**
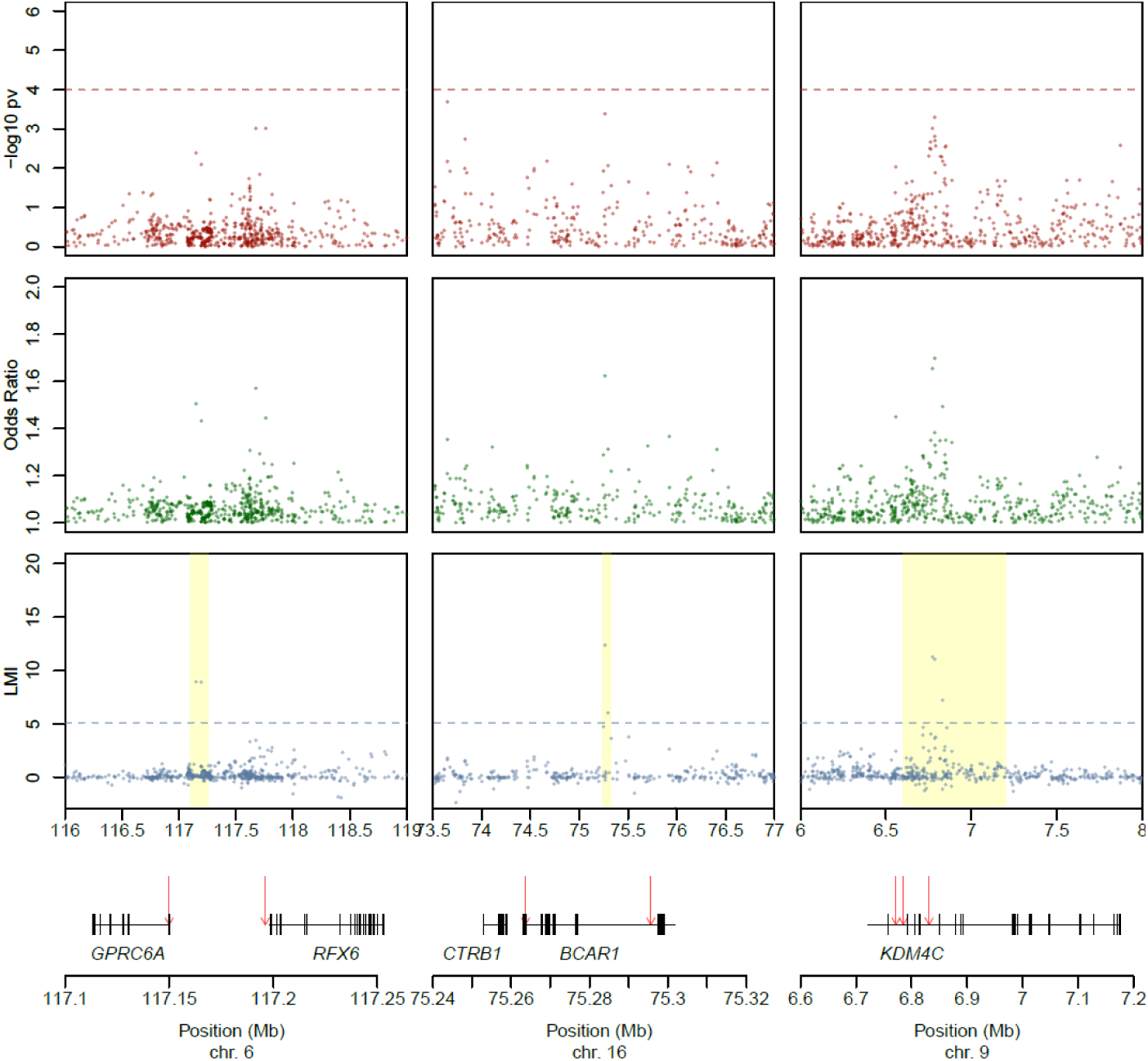
Scatterplots of the –log10 *p*-values, local Moran’s index (LMI) values and odds ratios (OR) for three genomic regions prioritized based on their LMI value. Highlighted regions show the hits identified in the 2D, but not in the 1D approach.

We also found 11 SNPs associated with *CDKN2A*, a gene that is almost universally inactivated in PC^32^ and that is mutated in some hereditary forms of PC^33,34^. Other SNPs identified by LMI were *DVL*-rs73185718 and *PRKCA*-rs11654719, that were also prioritized by GWAS, and two SNPs tagging *ROR2* (rs12002851 and rs2002478), a member of the *Wnt* pathway that plays a relevant role in PC^35^. *KDM4C-*rs72699638, a Lys demethylase 4C highly expressed in PC^36^ was also prioritized by LMI. Interestingly, the LMI analysis identified a hotspot region of 61 SNPs located upstream of *XBP1* (chr. 22, 28.3-29.3 Mb), a highly expressed gene in the healthy pancreas. The transcription factor XBP1 is involved in ER stress and the unfolded protein response - a highly relevant process in acinar homeostasis due to the high protein-producing capacity of these cells - and it plays an important role in pancreatic regeneration^37^.

#### Functionality of LMI-variants

We used CADD (Combined Annotation Dependent Depletion^38^ values to score the deleteriousness of LMI SNPs. LMI variant prioritization detected three variants in coding transcripts which showed the top CADD values and were not prioritized using the GWAS approach: *GPRC6A*-rs6907580 in chr6:117,150,008, CADD-score=5.0, LMI=8.93, GWAS *p-*value=4×10^−3^; *MS4A5*-rs34169848 in chr11:60,197,299, CADD-score=24.4, LMI=7.09, GWAS *p-*value=1×10^−2^; and *LRRC36*-rs8052655 in chr16:67,409,180, CADD-score=24.4, LMI=5.75, GWAS *p-*value=2×10^−2^. *GPRC6A*-rs6907580 is a well-characterized stop-gain variant in exon 1 of *GPRC6A* (*G protein-coupled receptor family C group 6 member A*). *GPRC6A* is expressed in pancreatic β-cells and participates in endocrine metabolism^39^ and this SNP is linked to a non-functional variant of *GPRC6A* receptor protein^40^. Furthermore, LMI identified rs17078438 (6q22.1) in *RFX6*, a pancreas-specific gene involved in endocrine development^41^ (**Figure 5**).

### 3D-Approach: genomic interaction analysis

To gain further insight into the putative biological functions of the 624 candidate SNPs selected through GWAS-LMI, we focused on a set of 6,761 significant chromatin interactions (*p-*values≤1×10^−5^) (see Methods) identified using Hi-C interaction pancreatic tissue maps at 40Kb resolution^42^. Throughout the rest of the text, we will refer to the chromatin interaction component containing the prioritized SNP as “bait” and to its interacting region as “target”. In total, 54 target regions overlapping with 37 genes interacted with bait regions harboring 76/624 (12.1%) SNPs (**Table S6**). Among them, we highlight again *XBP1* as we discovered that an intronic region of *TTC28* (bait: 22:28,602,352-28,642,352bp) including four LMI-selected SNPs (rs9620778, rs9625437, rs17487463 and rs75453968, all in high LD, *r*^*2*^>0.95, in CEU population) significantly interacted with the *XBP1* promoter (target: 22:29,197,371-29,237,371bp, *p-* value=1.3×10^−9^) (**Figure 6**). To confirm that this target region is relevant for pancreatic carcinogenesis, we retrieved from ENCODE the Chip-Seq data of all available non-tumor pancreatic samples (n=4 individuals) as well as from PANC-1 pancreas cancer cells (see Methods). We found that the H3K27Ac mark present in the *XBP1* promoter is completely lost in PANC-1 cells and is reduced in a sample of a Pancreatic Intraepithelial Neoplasia 1B, a PC precursor in comparison to normal pancreas (**Figure 6**). To characterize the bait and promoter regions upstream of *XBP1* further, we ran eight chromatin states using ChromHMM (**Supplementary Methods**). We observed a clear loss of enhancers/weak promoters in the corresponding target regions in the precursor lesions and in PANC-1 cells. This loss of activity is in line with the observation that *XBP1* expression is reduced in cancer. Moreover, small enhancers are also lost in the bait region of the aforementioned samples. We also checked whether the 3D maps for this region were comparable in healthy pancreas and PANC-1 cells and found that there was no significant contact in PANC-1 cells (**Figure 6**). Overall, these analyses indicate that the SNPs interacting in 3D space with the *XBP1* promoter could contribute to the differential expression of the gene associated with malignant transformation. These findings provide proof of concept that the LMI analysis combined with 3D genomics can contribute to decipher the biological relevance of orphan SNPs.

**Figure 6.**
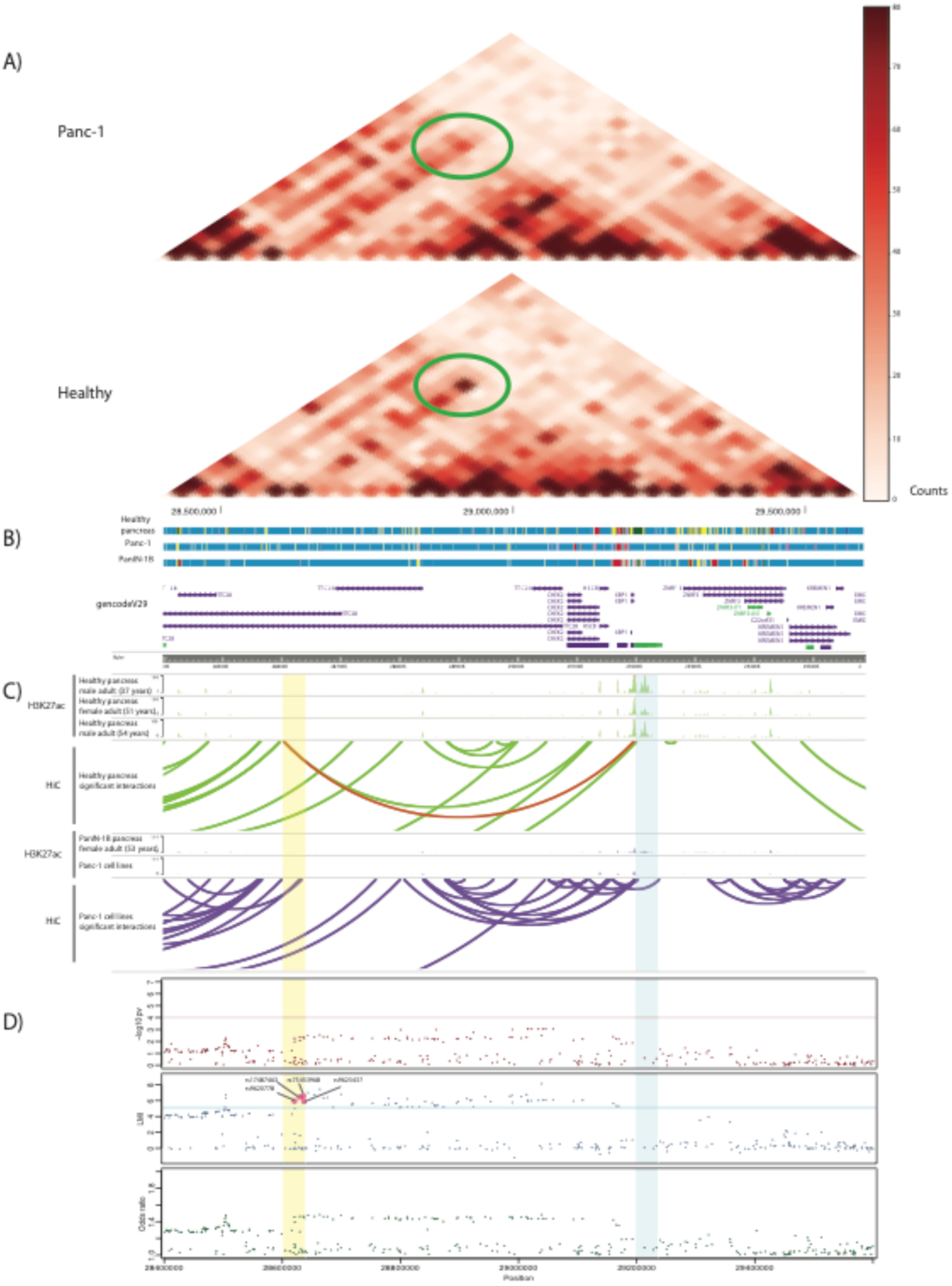
Three-dimensional genome organization in healthy and PANC-1 cells and association results corresponding to the genomic region around *XBP1* using the standard GWAS and 2D approaches. A) Coverage-normalized Hi-C maps of healthy samples and PANC-1 cells at 40Kb resolution. Green ellipses highlight the interaction between the region harboring four Local Moran’s Index (LMI)-selected SNPs and the *XBP1* promoter. B) Tracks of the ChromHMM Chromatin for 8 states in healthy pancreas, PANC-1 cells, and a Pancreatic Intraepithelial Neoplasia 1B. Promoters are colored in light purple, strong enhancers in dark green and weak enhancers in yellow. Note that the strong enhancer in the target region is lost in the PANC-1 and PanIN-1B samples, compared to the healthy samples. C) UCSC tracks of H3K27ac, an enhancer-associated mark, and arcs linking significant interactions called by Homer. Interactions in healthy pancreas samples are in green and those in PANC-1 and in the PanIN-1B sample are in purple. Red arc represents the interaction between LMI-prioritized SNPs and the *XBP1* promoter (highlighted region in Hi-C map in A). D) Scatterplots of SNPs in region chr22:28,400,000-29,600,000 (hg19) and their –log10 (p-value), LMI and odds ratio. Bait and target chromatin interaction regions are highlighted in yellow and blue, respectively.

To explore the translatable potential of the loci identified, we searched for all genes detected through GWAS-LMI and 3D genomic interactions in the PharmaGKB database (**Supplementary Material**). While we did not find direct evidence of these genes as targets for current PC treatments, 23/338 (6.8%) of the genes were annotated in the list of clinically actionable gene-drug associations for other cancer types or conditions associated with PC.

## DISCUSSION

In this work, we have expanded the scope of genomic analysis of the susceptibility to PC from the standard GWAS strategy to include novel approaches building on spatial autocorrelations of LMI and the 3D chromatin. An in-depth *in-silico* functional analysis leveraging available genomic information from public databases allowed us to prioritize novel candidate variants with strong biological plausibility. We have thus reached a novel landscape on the inherited basis of PC and have paved the way to the application of a similar strategy to any other human disease or interest.

This is the first PC GWAS involving an exclusively Europe-based population sample. Of the previously reported European ancestry population GWAS hits, 42.5% were replicated, supporting the methodological soundness of the study. The lack of replication of other PC GWAS hits may be explained by variation in the MAFs of the SNPs among Europeans, population heterogeneity, differences in the genotyping platform used, and differences in calling methods applied, among others. Replicated GWAS hits included *LINC00673*-rs7214041 reported to be in complete LD with *LINC00673-*rs11655237^11^, previously shown to be a PC-associated variant^9^ and replicated in our GWAS. *LINC00673* lies in a genomic region that is recurrently amplified and overexpressed in PC and is associated with poor clinical outcome^19^. Experimental evidence supports a functional role of *LINC00673* in the regulation of PC differentiation and in epithelial-mesenchymal transition^19^. Independent studies have confirmed the relevance of *LINC00673* in tumors and *in vitro*^43^. Beyond replicating previous GWAS hits, our study identified a novel variant in *NR5A2* (rs3790840) that independently associated with PC risk, strengthening the relevance of this gene in PC susceptibility.

We have also explored the potential of novel post-GWAS approaches to uncover variants failing to reach the strict GWAS *p*-value significance threshold. We applied the LMI for the first time in the genomics field (Anselin 1995). We replicated 6.4% of the previous reported GWAS Catalog signals for PC in European populations by considering the top 0.5% LMI variants, a LMI threshold that is overly conservative, given that many of the GWAS Catalog-replicated signals have lower LMI than the cut-off value we selected (see Methods). The ability of LMI to prioritize low MAF SNPs, unlike the GWAS approach, may also explain the low replicability rate. Despite the latter, LMI helps to identify correct signals within genomic regions, by scoring lower those regions that do not maintain LD structure (**Figure S5**).

To shed light into the functionality of the newly identified variants, we interrogated several databases at the SNP, gene, and pathway levels. We found sound evidence pointing to the functional relevance of several of the 143 GWAS *p*-value prioritized SNPs in the pancreas (**Table S3, Supplemental Material**). The importance of the multi-hit CASC8 region (8q24.21) is supported by post-GWAS *in-silico* functional analyses as well as by its previously associations with PC at the gene level^19^. In particular, 12/27 SNPs identified in *CASC8* were annotated as regulatory. Among them, *CASC8*-rs283705 and *CASC8*-rs2837237 (*r*^*2*^=0.68) are likely to be functional with a score of 2b in RegulomeDB (TF binding + any motif + DNase Footprint + DNase peak). Another variant (*CASC8*-rs1562430) was previously associated with risk of breast carcinoma^44^ and is in high LD (*r*^*2*^>0.85) with 18 *CASC8* prioritized variants. None of the prostate cancer-associated SNPs in *CASC8* overlapped with the 27 identified variants in our study. The fact that this gene has not been reported previously in other PC GWAS could be due to the different genetic background of the study populations or to an overrepresentation of the variants tagging *CASC8* in the Oncoarray platform used here.

In addition to confirming SNPs in *TERT*, we found strong evidence for the participation of novel susceptibility genes in telomere biology (*PARN*) and in the post-transcriptional regulation of gene expression (*PRKCA* and *EIF2B5*). Our study also expands the landscape of variants and genes involved in exocrine biology, including *SEC63, NOC2*/*RPH3AL* and *SCRT* whose function is likely to participate in acinar function and in acinar-ductal metaplasia, a PC pre-neoplastic lesion^45^.

The results from LMI and 3D genomic interactions further reinforce the role of genetic variation in these pathways. Among the SNP-LMI variants, *in-silico* functional assessment found evidence for a role of *GPRC6A*-rs6907580, *MS4A5*-rs34169848, *LRRC36*-rs8052655, *RFX6*-rs17078438, and *KDM4C*-rs72699638. The 3D genomic interaction approach also converged in *XBP1*, a critical regulator of acinar homeostasis. *XBP1* is a potential candidate detected through a previously uncharacterized “bait” SNP. These findings are particularly important considering that genetic mouse models have unequivocally shown that pancreatic ductal adenocarcinoma, the most common form of PC, can be initiated from acinar cells^46^. Similar results were found with other LMI selected SNPs associated with their target genes only by detecting significant spatial interactions between them (**Table S6**).

KEGG pathway enrichment analysis also validated other important pathways for PC, including “Glycosaminoglycan biosynthesis heparan sulfate” and “ERBB signaling pathway”. Heparan sulfate (HS) is formed by unbranched chains of disaccharide repeats which play roles in cancer initiation and progression^47^. Interestingly, the expression of HS proteoglycans increases in PC^48^ and related molecules, such as hyaluronic acid, are important therapeutic targets in PC^49,50^. ERBB signaling is important both in PC initiation and as a therapeutic target^51^.

The enrichment analysis indicates that urate levels, depression, and body mass index - three GWAS traits previously reported to be associated with PC risk - were enriched in our prioritized gene set. Urate levels have been associated with both PC risk and prognosis^52,53^. In addition, patients with lower relative levels of kynurenic acid have more depression symptoms^54^. PC is one of the cancers with the highest occurrence of depression preceding its diagnosis^55^. Furthermore, body mass index has been previously associated with PC risk in diverse populations^56-58^ and it has been suggested that increasing PC incidence may be partially attributed to the obesity epidemic. Insulin resistance is one of the mechanisms possibly underlying the obesity and PC association, through hyperinsulinemia and inflammation^59^.

Our post-GWAS approach has limitations that should be addressed in future studies. For example, our study has a relatively small sample size, some imbalances regarding gender and geographical areas, and the Hi-C maps that we used have limited resolution (40 kb). To account for population imbalances, regression models were adjusted for gender and for country of origin, as well as for first five principal components. In turn, our study has many strengths: a standardized methodology was applied in all participating centers to recruit cases and controls, to collect information, and to obtain and process biosamples; state-of-the-art methodology was used to extend the identification of variants, genes, and pathways involved in PC genetic susceptibility. Most importantly, the combination of GWAS, LMI and 3D genomics to identify new variants has not been applied in the past and has proven crucial to refine results, reduce the number of false positives, and establish whether borderline GWAS *p-*value signals could be true positives. These three strategies, together with an in-depth *in-*silico functional analysis, offer a comprehensive approach to advance the study of PC genetic susceptibility.

## METHODS

### 1D Approach: PanGenEU GWAS - *Single marker association analyses*

#### Study population

We used the resources from the PanGenEU case-control study conducted in Spain, Italy, Sweden, Germany, United Kingdom, and Ireland, between 2009-2014^60,61^. Eligible PC patients, men and women ≥18 years of age, were invited to participate. Eligible controls were hospital in-patients with primary diagnoses not associated with known risk factors of PC. Controls from Ireland and Sweden were population-based. Institutional review board approval and written informed consent was obtained from all participating centers and study participants, respectively. To increase statistical power, we included controls from the Spanish Bladder Cancer (SBC)/EPICURO study, carried out in the same geographical areas where PanGenEU Study was conducted. Characteristics of the study populations are detailed in **Table S7**.

#### Genotyping and quality control in the PanGenEU study

DNA samples were genotyped using the Infinium OncoArray-500K at the CEGEN facility (Spanish National Cancer Research Centre, CNIO). Genotypes were called using GenTrain 2.0 cluster algorithm in GenomeStudio software v.2011.1.0.24550 (Illumina, San Diego, CA). Genotyping quality control criteria considered the missing call rate, unexpected heterozygosity, discordance between reported and genotyped gender, unexpected relatedness, and estimated European ancestry <80%. After removing samples that did not pass the quality control filters, duplicated samples, and individuals with incomplete data regarding age of diagnosis/recruitment, 1,317 cases and 700 controls were available for the association analyses. SNPs in sex chromosomes and those that did not pass the Hardy-Weinberg equilibrium (*p*-value<10^−6^) were also discarded. Overall, 451,883 SNPs passed the quality control filters conducted before the imputation.

#### Genotyping and quality control of SBC/EPICURO controls

Genotyping of germline DNA was performed using the Illumina 1M Infinium array at the NCI Core Genotyping Facility as previously described^62^, which provided calls for 1,072,820 SNP genotypes. We excluded SNPs in sex chromosomes, those with a low genotyping rate (<95%), and those that did not pass the Hardy-Weinberg equilibrium threshold. In addition, the exome of 36 controls was sequenced with the TruSeq DNA Exome and a standard quality control procedure both at the SNP and individual level was applied: SNPs with read depth <10 and those that did not pass the tests of base sequencing quality, strand bias or tail distance bias, were considered as missing and imputed (see *Imputation* section for further details). Overall, 1,122,335 SNPs were available for imputation. In total, 916 additional controls were considered for this analysis.

#### Imputation

Imputation was performed at the Wellcome Sanger Institute, Cambridge, UK, and CNIO, Madrid, Spain, for the PanGenEU and the SBC/EPICURO studies, respectively. Imputation of missing genotypes was performed using IMPUTE v2^63^ and genotypes of SBC/EPICURO controls were pre-phased to produce best-guess haplotypes using SHAPEIT v2 software^64^. For both PanGenEU and EPICURO studies, the 1000 G (Phase 3, v1) reference data set was used^17^.

#### Association analyses

A final set of 317,270 common SNPs (MAF>0.05) that passed quality control in both studies and showed comparable MAF across genotyping platforms was considered for analysis. We ensured the inclusion of the 40 variants previously associated with PC risk in individuals of Caucasian origin compiled in GWAS Catalog^5^. Logistic regression models were computed assuming an additive mode of inheritance for the SNPs, adjusted for age at PC diagnosis or at control recruitment, sex, the area of residence [Northern Europe (Germany and Sweden), European islands (UK and Ireland), and Southern Europe (Italy and Spain)], and the first 5 principal components (PCs) calculated with *prcomp* R function based on the genotypes of 32,651 independent SNPs, (J Tyrer, personal communication) to control for potential population substructure.

#### Validation of the novel GWAS hits

To replicate the top 20 associations identified in the Discovery phase, we performed a meta-analysis using risk estimates obtained in previous GWAS studies from the Pancreatic Cancer Cohort Consortium (PanScan: https://epi.grants.cancer.gov/PanScan/) and the Pancreatic Cancer Case-Control Consortium (PanC4: http://www.panc4.org/), based on 16 cohort and 13 case-control studies. Details on individual studies, namely PanScan I, PanScan II, PanScan III and PanC4, have been described elsewhere^6-9^. Genotyping for PanScan studies was performed at the Cancer Genomic Research Laboratory (CGR) of the National Cancer Institute (NCI) using HumanHap550v3.0, and Human 610-Quad genotyping platforms for PanScan I and II, respectively, and the Illumina Omni series arrays for PanScan III. Genotyping for PanC4 was performed at the Johns Hopkins Center for Inherited Disease Research using the Illumina HumanOmniExpressExome-8v1 array. PanScan I/II datasets were imputed together using the 1000 G (Phase3, v1) reference data set^17^ and IMPUTE2^63^ and adjusting for study (PanScan I and II), geographical region (for PanScan III), age, sex, and PCA of population substructure (5 PC’s for PanScan I+II, 6 for PanScan III) for PanScan models, and for study, age, sex and 7 PCA population substructure for PanC4 models. Summary statistics from PanScanI/II, PanScan III and PanC4 were used for a meta-analysis using a random-effects model based on effect estimates and standard errors with the metafor R package^65^.

#### Post-GWAS functional *in silico* analysis

An exhaustive *in-silico* analysis was conducted for associations with *p*-values<1×10^−4^ in the PanGenEU GWAS (**Figure S3**). Bioinformatics assessments included evidence of functional impact^66,67^, annotation in overlapping genes and pathways^66^, methylation quantitative trait locus in leukocyte DNA from a subset of the PanGenEU controls (mQTLs), expression QTL (eQTLs) in normal and tumoral pancreas (GTEx and TCGA, respectively)^68,69^, annotation in PC-associated long non-coding RNA (lncRNAs)^19^, protein quantitative trait locus analysis in plasma (pQTLs)^70^, overlap with regulatory chromatin marks in pancreatic tissue obtained from ENCODE^71^, association with relevant human diseases^72^, and annotation in differentially open chromatin regions (DORs) in human pancreatic cells^41^. We also investigated whether prioritized variants had been previously associated with PC comorbidities or other types of cancers^5^. Furthermore, we used HOMER to map SNPs to significant 3D chromatin interaction (CI) in healthy pancreas tissue^26^. Then, we annotated those SNPs in significant interaction regions with the chromatin states^73^.

In addition to the functional analyses at the variant level, we conducted enrichment analyses at the gene level using the FUMAGWAS web tool^72^ and investigated whether our prioritized set of genes appeared altered at the tumor level in a collection of pancreatic tumor samples^74^. Methodological details of all bioinformatics analyses conducted are described in detail in Supplementary Material.

### 2D Approach: Local Moran Index

#### Local Moran’s Index calculation

The LMI was obtained for each SNP considered in the GWAS (n=317,270) using the summary statistics resulting from the association analyses as follows. First, the OR of each SNP was referred to its risk-increasing allele (*i.e.*, OR>1) and the distribution of ORs was transformed to the inverse of the normal distribution. Second, each SNP was matched by MAF with surrounding common SNPs (*i.e.*, SNPs with MAF>=1% in the 85 European individuals of the 1000G, Phase 1, version 3), considering a window of +/- 500kb to ensure that haplotypes were matched. Linkage disequilibrium (*r*^*2*^) was used as a proxy for the distance between each SNP and each of its neighboring SNPs. Next, the LMI for *i*-th SNP was calculated as:

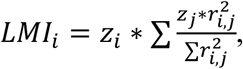

where *LMI*_*i*_ is the LMI value for the *i*-th SNP; *z*_*i*_ is the OR value for the *i*-th SNP, obtained from the inverse of the normal distribution of ORs for all SNPs; *z*_*j*_ is the OR for the *j-*th SNP within the physical distance and MAF-matched defined bounds; and 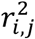 is the LD value, measured by *r*^*2*^, between the *i-*th SNP and the *j-*th SNP.

After LMI calculation for the full set of SNPs, we discarded the SNPs that: (1) had a negative LMI, meaning either that surrounding SNPs and target SNP have largely different ORs or that they are in linkage equilibrium and, therefore, do not pertain to the same cluster; or (2) had a positive LMI, i.e. target and surrounding SNPs have similar ORs, but the SNP came from the bottom 50% tail of the distribution of the ordered transformed OR distribution. This generated a total final set of 102,146 SNPs, out of which we selected the top 0.5%, at a threshold of LMI value = 5.1071 (n=510).

To assess the usefulness of the LMI score for SNP prioritization, we ran two tests using SNPs known to be associated with PC in European populations [GWAS Catalog, n=40^5^]. Before performing this benchmarking test, we corrected the 40 signals by LD using a custom made “*greedy*” algorithm. First, we calculated all pairwise LD values (*r*^2^) for all the SNPs on the same chromosome. Then, we reviewed the list of SNPs ordered by ascending position chromosome-wise and considered as a cluster all the SNPs that had *r*^*2*^>0.2 with the SNP under consideration. We considered this set of SNPs as a unique genomic signal, filtered out the SNPs assigned to the cluster from the ranked list, and then proceeded to the next SNP. This resulted in a total of 30 independent clusters of ≥1 SNPs. When more than one SNP was included within the same cluster, the SNP with the highest LMI was selected. The same procedure was applied to identify the independent loci in the GWAS-selected SNPs (n=143) and the set of LMI-selected SNPs (n=624).

For the first benchmarking test, we first evaluated whether the GWAS Catalog PC-associated SNPs had a LMI value higher than expected. Then, we ranked the LMI value for the 102,146 LMI-selected SNPs from highest to lowest, assigning position number “1” to the SNP with the highest LMI (LMI=18.23) and position number “102,146” to the one with the lowest LMI (LMI=0.000001). Out of the 30 signals derived from the GWAS Catalog, 22 were present in our 102,146 selected set. The observed median rank position in this list for the 22 PC signals was 22,640. This average position was significantly higher than 10,000 randomly selected sets of the same size (one tail *p-* value=0.0013) (**Figure S6**). Loci annotated in the GWAS Catalog as associated with PC tend to score higher LMI than expected by chance. Finally, for the benchmark based on replication of loci, out of the 30 independent signals, 21 clusters of more than one SNP were considered as replicated signals and 9 SNPs that were found by only one study were not replicated.

#### Biological annotation and functional in-silico analysis

LMI-selected variants were annotated to genes using annotatePeaks.pl script in HOMER^26^ and their functionality was predicted using CADD^38^ online software.

### 3D Approach: Hi-C pancreas interaction maps and interaction selection

The 3D Hi-C interaction maps for both healthy pancreas tissue (Schmitt et al. 2016) and for a pancreatic cancer cell line (PANC-1) were generated using TADbit as previously described^75^. Briefly, Hi-C FASTQ files for 7 replicas of healthy pancreas tissue were downloaded from GEO repository (Accession number: GSE87112; Sequence Read Archive Run IDs: SRR4272011, SRR4272012, SRR4272013, SRR4272014, SRR4272015, SRR4272016, SRR4272017) and for PANC-1 FASTQ, files were available from ENCODE (Accession number: ENCSR440CTR). For further analysis, all 7 healthy samples were merged. Next, the FASTQ files were mapped against the human reference genome hg19, parsed and filtered with TADbit to get the final number of valid interacting read pairs. Total numbers of 99,074,082 and 287,201,883 valid interaction pairs were obtained for the healthy and PANC-1 datasets, respectively. Valid pairs were next binned at 40 kb resolution to obtain chromosome-wide interaction matrices. Next, the HOMER package^26^ was used to detect significant interactions between two bins of 40kb within a window of 4Mb using the –center and --maxDist 2000000 parameters. Using HOMER’s default parameters (significant interactions at *p*-value=0.001), the final number of nominally significant interactions was 41,833 for the healthy dataset and 357,749 for the PANC-1 dataset. To further filter the interactions, we assessed the number of possible unique bin combinations within 2Mb of a bin (that is, 4,950 combinations of any two bins) and calculated those interactions that passed a Bonferroni corrected threshold *p*-value=10^−5^. The sub-selected set of interactions was reduced to 6,761 for the healthy sample (that is, 16.2% top interactions from those originally selected by HOMER default parameters). Next, we sub-sampled the top 16% interactions for PANC-1 list, resulting in 57,813 significant interactions.

## ACKNOWLEDGEMENTS

The authors are thankful to the patients, coordinators, field and administrative workers, and technicians of the European Study into Digestive Illnesses and Genetics (PanGenEU) and the Spanish Bladder Cancer (SBC/EPICURO) studies. We also thank Marta Rava, former member of the GMEG-CNIO, Guillermo Pita and Anna González-Neira from CGEN-CNIO, and Joe Dennis and Laura Fachal from the University of Cambridge, for genotyping PanGenEU samples, performing variant calling and SNP imputation, and editing data.

## AUTHOR CONTRIBUTIONS

Study conception: **NM, ELM.**

Design of the work**: ELM, JAR, DE, MAMR, FXR.**

Data acquisition**: EMM, PGR, RTL, AC, MH, MI, XM, ML, CWM, JP, MOR, BMB, AT, AF, LMB, TCJ, LDM, TG, WG, LS, LA, LC, JB, EC, LI, JK, NK, MM, JM, DOD, AS, WY, JY, PanGenEU Investigators, MGC, MK, NR, DS, SBC/EPICURO Investigators, DA, AAA, LBF, PMB, PB, BBM, JB, FC, MD, SG, JMG, PJG, MG, LLM, LD, NRN, UP, GMP, HAR, MJS, XOS, LDT, KV, WZ, SC, BMW, RZSS, APK, LA, FXR, NM.**

Data analysis**: ELM, JAR, LA, OL.**

Interpretation of data**: ELM, JAR, LA, OL, EMM, MAMR, FXR, NM.**

Creation of new software used in the work**: ELM, LA, JAR, OL, MAMR.**

To have drafted the work or substantively revised it**: ELM, JAR, LA, OL, TCJ, UP, HAR, APK, LA, MAMR, FXR, NM.**

To have approved the submitted version (and any substantially modified version that involves the author’s contribution to the study): **ALL AUTHORS**

To have agreed both to be personally accountable for the author’s own contributions and to ensure that questions related to the accuracy or integrity of any part of the work, even ones in which the author was not personally involved, are appropriately investigated, resolved, and the resolution documented in the literature: **ALL AUTHORS**.

## REFERENCES

1. Carrato, A. et al. A Systematic Review of the Burden of Pancreatic Cancer in Europe: Real-World Impact on Survival, Quality of Life and Costs.” J Gastrointest Cancer 46, 201–11 (2015).

2. Malvezzi, M. et al. European Cancer Mortality Predictions for the Year 2015: Does Lung Cancer Have the Highest Death Rate in EU Women?” Ann Oncol 26, 779–86 (2015).

3. Torre, L. A., Siegel, R. L. Ward, E. M. & Jemal, A. Global Cancer Incidence and Mortality Rates and Trends - An Update. Cancer Epidemiol Biomarkers Prev 25, 16–27 (2016).

4. Rahib, L. et al. Projecting Cancer Incidence and Deaths to 2030: The Unexpected Burden of Thyroid, Liver, and Pancreas Cancers in the United States. Cancer Res 74, 2913–21 (2014).

5. Buniello, A. et al. The NHGRI-EBI GWAS Catalog of Published Genome-Wide Association Studies, Targeted Arrays and Summary Statistics 2019. Nucleic Acids Res 47, D1005–12 (2019).

6. Amundadottir, L. et al. Genome-Wide Association Study Identifies Variants in the ABO Locus Associated with Susceptibility to Pancreatic Cancer. Nature Genet 41, 986–90 (2009).

7. Petersen, G. M. et al. A Genome-Wide Association Study Identifies Pancreatic Cancer Susceptibility Loci on Chromosomes 13q22.1, 1q32.1 and 5p15.33. Nat Genet 42, 224–28 (2010).

8. Wolpin, B. M. et al. Genome-Wide Association Study Identifies Multiple Susceptibility Loci for Pancreatic Cancer. Nat Genet 46 994–1000 (2014).

9. Childs, E. J. et al. Common Variation at 2p13.3, 3q29, 7p13 and 17q25.1 Associated with Susceptibility to Pancreatic Cancer. Nature Genet 47, 911–16 (2015).

10. Zhang, M. et al. Three New Pancreatic Cancer Susceptibility Signals Identified on Chromosomes 1q32.1, 5p15.33 and 8q24.21. Oncotarget 7, 66328–43 (2016).

11. Klein, A. P. et al. Genome-Wide Meta-Analysis Identifies Five New Susceptibility Loci for Pancreatic Cancer. Nat Commun 9, 556 (2018).

12. Chen, F. et al. Analysis of Heritability and Genetic Architecture of Pancreatic Cancer: A PANC4 Study. Cancer Epidemiol Biomarkers Prev 28, 1238–45 (2019).

13. Anselin, L. Local Indicators of Spatial Association—LISA. Geograph Anal 27, 93–115 (1995).

14. Dekker, J., Rippe, K., Dekker, M., & Kleckner, N. Capturing Chromosome Conformation. Science 295, 1306–11 (2002).

15. Claussnitzer, M. et al. FTO Obesity Variant Circuitry and Adipocyte Browning in Humans. New Engl J Med 373, 895–907 (2015).

16. Montefiori, L. E. et al. A Promoter Interaction Map for Cardiovascular Disease Genetics. ELife 7, e35788 (2018).

17. Altshuler, D. L. et al. A Map of Human Genome Variation from Population-Scale Sequencing. Nature 467, 1061–73 (2010).

18. Li, D. et al. Pathway Analysis of Genome-Wide Association Study Data Highlights Pancreatic Development Genes as Susceptibility Factors for Pancreatic Cancer. Carcinogenesis 33, 1384–90 (2012).

19. Arnes, L. et al. Comprehensive Characterisation of Compartment-Specific Long Non-Coding RNAs Associated with Pancreatic Ductal Adenocarcinoma. Gut 68, 499–511 (2019).

20. Linxweiler, M, Schick, B. & Zimmermann, R. Let’s Talk about Secs: Sec61, Sec62 and Sec63 in Signal Transduction, Oncology and Personalized Medicine. Signal Transduct Target Ther 2:17002 (2017).

21. Matsumoto, M. et al. Noc2 Is Essential in Normal Regulation of Exocytosis in Endocrine and Exocrine Cells. Proc Natl Acad Sciences USA 101, 8313–18 (2004).

22. Risch HA. Etiology of pancreatic cancer, with a hypothesis concerning the role of N-nitroso compounds and excess gastric acidity. J Natl Cancer Inst 95, 948–60 (2003).

23. Körner, M. et al. Secretin Receptors in Normal and Diseased Human Pancreas: Marked Reduction of Receptor Binding in Ductal Neoplasia. Am J Pathol 167, 959–68 (2005).

24. Pemberton, J. R. Retention of Mercurial Preservatives in Desiccated Biological Products. J Clin Microbiol 2, 549–51 (1975).

25. Cobo, I. et al. Transcriptional Regulation by NR5A2 Links Differentiation and Inflammation in the Pancreas. Nature 554, 533–37 (2018).

26. Heinz, S. et al. Simple Combinations of Lineage-Determining Transcription Factors Prime Cis-Regulatory Elements Required for Macrophage and B Cell Identities. Mol Cell 38, 576–89 (2010).

27. Duan, B. et al. Genetic Variants in the Platelet-Derived Growth Factor Subunit B Gene Associated with Pancreatic Cancer Risk. Int J Cancer 142, 1322–31 (2018).

28. Rosendahl, J. et al. Genome-Wide Association Study Identifies Inversion in the CTRB1-CTRB2 Locus to Modify Risk for Alcoholic and Non-Alcoholic Chronic Pancreatitis. Gut 67, 1855–63 (2018).

29. Morris, A. P. et al. Large-Scale Association Analysis Provides Insights into the Genetic Architecture and Pathophysiology of Type 2 Diabetes. Nat Genet 44, 981–90 (2012).

30. Xue, A. et al. Genome-Wide Association Analyses Identify 143 Risk Variants and Putative Regulatory Mechanisms for Type 2 Diabetes. Nat Commun 9 2941 (2018).

31. Tang, Z. et al. GEPIA: A Web Server for Cancer and Normal Gene Expression Profiling and Interactive Analyses. Nucleic Acids Res 45, W98–102 (2017).

32. Notta, F. et al. A Renewed Model of Pancreatic Cancer Evolution Based on Genomic Rearrangement Patterns. Nature 538, 378–82 (2016).

33. Bartsch, D. K. et al. CDKN2A Germline Mutations in Familial Pancreatic Cancer. Ann Surg 236, 730–37 (2002).

34. Lynch, H. T. et al. Phenotypic Variation in Eight Extended CDKN2A Germline Mutation Familial Atypical Multiple Mole Melanoma-Pancreatic Carcinoma-Prone Families: The Familial Atypical Multiple Mole Melanoma-Pancreatic Carcinoma Syndrome. Cancer 94, 84–96 (2002).

35. Morris, J. P., Wang, S. C. & Hebrok, M. KRAS, Hedgehog, Wnt and the Twisted Developmental Biology of Pancreatic Ductal Adenocarcinoma. Nat Rev Cancer 10, 693–95 (2010).

36. Gregory, B. L. & Cheung, V. G. Natural Variation in the Histone Demethylase, KDM4C, Influences Expression Levels of Specific Genes Including Those That Affect Cell Growth. Genome Res 24, 52–63 (2014).

37. Hess, D. A. et al. Extensive Pancreas Regeneration Following Acinar-Specific Disruption of Xbp1 in Mice. Gastroenterology 141, 1463–72 (2011).

38. Rentzsch, P., Witten, D., Cooper, G. M., Shendure, J. & Kircher, M. CADD: Predicting the Deleteriousness of Variants throughout the Human Genome. Nucleic Acids Res 47, D886–94 (2019).

39. Pi, M. & Quarles, L. D. Multiligand Specificity and Wide Tissue Expression of GPRC6A Reveals New Endocrine Networks. Endocrinology (2012).

40. Jørgensen, S. et al. Genetic Variations in the Human g Protein-Coupled Receptor Class C, Group 6, Member A (GPRC6A) Control Cell Surface Expression and Function. J Biol Chem 292, 1524–34 (2017).

41. Arda, H. et al. A Chromatin Basis for Cell Lineage and Disease Risk in the Human Pancreas. Cell Syst 7, 310-322.e4 (2018).

42. Schmitt, A. D. et al. A Compendium of Chromatin Contact Maps Reveals Spatially Active Regions in the Human Genome. Cell Rep 17, 2042–59 (2016).

43. Zhao, X. et al. Linc00511 Acts as a Competing Endogenous RNA to Regulate VEGFA Expression through Sponging Hsa-MiR-29b-3p in Pancreatic Ductal Adenocarcinoma. J Cell Mol Med 22, 655–67 (2018).

44. Turnbull, C. et al. Genome-Wide Association Study Identifies Five New Breast Cancer Susceptibility Loci. Nat Genet 42, 504–7 (2010).

45. Notta, F., Hahn, S.A. & Real, F.X. The genetic roadmap of pancreatic cancer: still evolving. Gut 66, 2170–278 (2017).

46. Guerra, C. et al. Chronic Pancreatitis Is Essential for Induction of Pancreatic Ductal Adenocarcinoma by K-Ras Oncogenes in Adult Mice. Cancer Cell 11, 291–302 (2007).

47. Nagarajan, A., Malvi, P. & Wajapeyee, N. Heparan Sulfate and Heparan Sulfate Proteoglycans in Cancer Initiation and Progression. Front Endocrinol 9, 483 (2018).

48. Theocharis, A. D., Skandalis, S. S., Tzanakakis, G. N. & Karamanos, N. K. Proteoglycans in Health and Disease: Novel Roles for Proteoglycans in Malignancy and Their Pharmacological Targeting. FEBS J 277, 3904–23 (2010).

49. Hingorani, S. R. et al. HALO 202: Randomized Phase II Study of PEGPH20 Plus Nab-Paclitaxel/Gemcitabine Versus Nab-Paclitaxel/Gemcitabine in Patients With Untreated, Metastatic Pancreatic Ductal Adenocarcinoma. J Clin Oncol 36, 359–66 (2018).

50. Provenzano, P. P. et al. Enzymatic Targeting of the Stroma Ablates Physical Barriers to Treatment of Pancreatic Ductal Adenocarcinoma. Cancer Cell 21, 418–29 (2012).

51. Mccleary-Wheeler, A. L., Mcwilliams, R. & Fernandez-Zapico, M. E. Aberrant Signaling Pathways in Pancreatic Cancer: A Two Compartment View. Mol Carcinog 51, 25–39 (2012)

52. Mayers, J. R. et al. Elevation of Circulating Branched-Chain Amino Acids Is an Early Event in Human Pancreatic Adenocarcinoma Development. Nat Med 20, 1193–98 (2014).

53. Stotz, M. et al. Evaluation of Uric Acid as a Prognostic Blood-Based Marker in a Large Cohort of Pancreatic Cancer Patients. PLoS ONE 9, e104730 (2014).

54. Botwinick, I. C. et al. A Biological Basis for Depression in Pancreatic Cancer. Hpb (Oxford) 16, 740–43 (2014).

55. Eguia, V., Gonda T. A. & Saif MW. Early Detection of Pancreatic Cancer. JOP 13, 131–4 (2012).

56. Carreras-Torres, R. et al. The Role of Obesity, Type 2 Diabetes, and Metabolic Factors in Pancreatic Cancer: A Mendelian Randomization Study.” J Natl Cancer Inst 109 (2017).

57. Koyanagi, Y. N. et al. Body-Mass Index and Pancreatic Cancer Incidence: A Pooled Analysis of Nine Population-Based Cohort Studies with More than 340,000 Japanese Subjects. J Epidemiol 28, 245–52 (2018).

58. Lauby-Secretan, B. et al. Body Fatness and Cancer - Viewpoint of the IARC Working Group. New Engl J Med 375, 794–98 (2016).

59. Park, J., Morley, T. S., Kim, M., Clegg, D. J. & Scherer, D. J. Obesity and Cancer -Mechanisms Underlying Tumour Progression and Recurrence. Nat Rev Endocrinol 10, 455–65 (2014).

60. Gomez-Rubio, P. et al. Reduced risk of pancreatic cancer associated with asthma and nasal allergies. Gut 66, 314–322 (2017).

61. Molina-Montes, E. et al. Risk of pancreatic cancer associated with family history of cancer and other medical conditions by accounting for smoking among relatives. Int J Epidemiol 47, 473–483 (2018).

62. Rothman, N. et al. A Multi-Stage Genome-Wide Association Study of Bladder Cancer Identifies Multiple Susceptibility Loci. Nat Genet 42, 978–84 (2010).

63. Howie, B. N., Donnelly, P. & Marchini, J. A Flexible and Accurate Genotype Imputation Method for the next Generation of Genome-Wide Association Studies. PLoS Genet 5, e1000529 (2009).

64. Delaneau, O., Marchini, J. & Zagury, J. F. A Linear Complexity Phasing Method for Thousands of Genomes.” Nat Methods 9, 179–81 (2012).

65. Viechtbauer, W. Conducting Meta-Analyses in R with the Metafor. J Stat Software 36, 1–48 (2010).

66. McLaren, W. et al. The Ensembl Variant Effect Predictor. Genome Biol 17, 122 (2016).

67. Martín-Antoniano, I., Alonso, A., Madrid, M., López De Maturana, E. & Malats, N. DoriTool: A Bioinformatics Integrative Tool for Post-Association Functional Annotation. Public Health Genomics 20, 126–35 (2017).

68. Ardlie, K. G. et al. The Genotype-Tissue Expression (GTEx) Pilot Analysis: Multitissue Gene Regulation in Humans. Science 348, 648–60 (2015).

69. Gong, J. et al. PancanQTL: Systematic Identification of Cis -EQTLs and Trans - EQTLs in 33 Cancer Types.” Nucleic Acids Res 46, D971–76 (2018).

70. Sun, B. B. et al. Genomic Atlas of the Human Plasma Proteome. Nature 558, 73–79 (2018).

71. Sloan, C. A. et al. ENCODE Data at the ENCODE Portal. Nucleic Acids Res 44 D726–32 (2016).

72. Watanabe, K., Taskesen, E., Van Bochoven, A. & Posthuma, D. Functional Mapping and Annotation of Genetic Associations with FUMA. Nat Commun 8, 1826 (2017).

73. Ernst, J. & Kellis, M. ChromHMM: Automating Chromatin-State Discovery and Characterization. Nat Methods 9, 215–6 (2012).

74. Gao, J. et al. Integrative Analysis of Complex Cancer Genomics and Clinical Profiles Using the CBioPortal.” Sci Signaling 6, pl1 (2013).

75. Serra, F. et al. Automatic Analysis and 3D-Modelling of Hi-C Data Using TADbit Reveals Structural Features of the Fly Chromatin Colors. PLoS Comput Biol 13, e1005665 (2017).

