## Supplementary Material for "A multilayered post-GWAS assessment on genetic susceptibility to pancreatic cancer"

**Annex 1.** PanGenEU Investigators.

**Annex 1.** SBC/EPICURO Investigators.

Supplementary Methods and Results.

**Table S1.** Replication of the SNPs reported as associated with PC risk in European population and published in GWAS Catalog (.docx file).

**Table S2**. Validated variants (at the nominal p-value) in PanScan and PanC4 populations, among the top 20 SNPs identified in the PanGenEU GWAS study (.docx file).

**Table S3**. Annotation and functional *in silico* analysis of the 143 prioritized SNPs (.xls file).

**Table S4.** Results from the gene enrichment analysis performed with FUMA (.txt file).

**Table S5.** List of the 624 SNPs prioritized according to their local Moran index (LMI) (.xls file).

**Table S6.** List of the 76 SNPs overlapping with a chromatin interaction region (bait regions) and their 54 targets (.xls file).

**Table S7.** Characteristics of the study populations (.docx file).

**Figure S1**. GWAS Manhattan plot for the PanGenEU study.

**Figure S2**. Q-Q plots for the PanGenEU (S2a) and PanGenEU&EPICURO study populations (S2b).

**Figure S3**. Functional *in silico* analysis strategy.

**Figure S4**. Complementary NETWORK (in blue our input pathways; in green, the complementary nodes included) obtained with Pathway-connector webtool.

**Figure S5**. Minor allele frequency (MAF) distributions for the top 97 SNPs identified by LMI (in pink) and by GWAS (in blue).

**Figure S6**. Results of the benchmarking test showing that the median rank position of the LMI values for the 22 PC signals from the GWAS Catalog is significantly higher than 10,000 randomly selected sets of the same size.

### Annex 1. PanGenEU Centres and Investigators

Spanish National Cancer Research Centre (CNIO), Madrid, Spain: Núria Malats[1], Francisco X Real[1], Evangelina López de Maturana, Paulina Gómez-Rubio, Esther Molina-Montes, Lola Alonso, Mirari Márquez, Roger Milne, Ana Alfaro, Tania Lobato, Lidia Estudillo.

Verona University, Italy: Rita Lawlor[1], Aldo Scarpa, Stefania Beghelli.

National Cancer Registry Ireland, Cork, Ireland: Linda Sharp[1], Damian O'Driscoll.

Hospital Madrid-Norte-Sanchinarro, Madrid, Spain: Manuel Hidalgo[1], Jesús Rodríguez Pascual.

Hospital Ramon y Cajal, Madrid, Spain: Alfredo Carrato[1], Carmen Guillén-Ponce, Mercedes Rodríguez-Garrote, Federico Longo-Muñoz, Reyes Ferreiro, Vanessa Pachón, M Ángeles Vaz.

Hospital del Mar, Barcelona, Spain: Lucas Ilzarbe[1], Cristina Álvarez-Urturi, Xavier Bessa, Felipe Bory, Lucía Márquez, Ignasi Poves✝, Fernando Burdío, Luis Grande, Mar Iglesias, Javier Gimeno.

Hospital Vall d´Hebron, Barcelona, Spain: Xavier Molero[1], Luisa Guarner✝, Joaquin Balcells.

Technical University of Munich, Germany: Christoph Michalski[1], Jörg Kleeff, Bo Kong.

Karolinska Institute, Stockholm, Sweden: Matthias Löhr[1], Jiaqui Huang, Weimin Ye, Jingru Yu.

Hospital 12 de Octubre, Madrid, Spain: José Perea[1], Pablo Peláez.

Hospital de la Santa Creu i Sant Pau, Barcelona, Spain: Antoni Farré[1], Josefina Mora, Marta Martín, Vicenç Artigas, Carlos Guarner, Francesc J Sancho, Mar Concepción, Teresa Ramón y Cajal.

The Royal Liverpool University Hospital, UK: William Greenhalf[1], Eithne Costello.

Queen's University Belfast, UK: Michael O'Rorke[1], Liam Murray✝, Marie Cantwell.

Laboratorio de Genética Molecular, Hospital General Universitario de Elche, Spain: Víctor M Barberá[1], Javier Gallego.

Instituto Universitario de Oncología del Principado de Asturias, Oviedo, Spain: Adonina Tardón[1], Luis Barneo.

Hospital Clínico Universitario de Santiago de Compostela, Spain: Enrique Domínguez Muñoz[1], Antonio Lozano, Maria Luaces.

Hospital Clínico Universitario de Salamanca, Spain: Luís Muñoz-Bellvís[1], J.M.

Sayagués Manzano, M.L. Gutíerrrez Troncoso, A. Orfao de Matos.

University of Marburg, Department of Gastroenterology, Phillips University of Marburg, Germany: Thomas Gress[1], Malte Buchholz, Albrecht Neesse.

Queen Mary University of London, UK: Tatjana Crnogorac-Jurcevic[1], Hemant M Kocher, Satyajit Bhattacharya, Ajit T Abraham, Darren Ennis, Thomas Dowe, Tomasz Radon

Scientific advisors of the PanGenEU Study: Debra T Silverman (NCI, USA) and Douglas Easton (U. of Cambridge, UK)

---

[1] Principal Investigator in each centre

**Annex 2. Spanish Bladder Cancer/EPICURO Centres and Investigators**

Institut Municipal d'Investigació Mèdica, Universitat Pompeu Fabra, Barcelona –Coordinating Center1: M. Kogevinas, N. Malats, F.X. Real, M. Sala, G. Castaño, M. Torà, D. Puente, C. Villanueva, C. Murta-Nascimento, J. Fortuny, E. López, S. Hernández, R. Jaramillo, G. Vellalta, L. Palencia, F. Fermández, A. Amorós, A. Alfaro, G. Carretero.

National Cancer Institute, NIH, USA –Coordinating Center2: Debra Silverman, Mustafa Dosemeci✝, Nathaniel Rothman, Montserrat García-Closas, Stephen Chanock.

Hospital del Mar, Universitat Autònoma de Barcelona, Barcelona: J. Lloreta, S. Serrano, L. Ferrer, A. Gelabert, J. Carles, O. Bielsa, K. Villadiego.

Hospital Germans Trias i Pujol, Badalona, Barcelona: L. Cecchini, J.M. Saladié, L. Ibarz.

Hospital de Sant Boi, Sant Boi de Llobregat, Barcelona: M. Céspedes.

Consorci Hospitalari Parc Taulí, Sabadell: C. Serra, D. García, J. Pujadas, R. Hernando, A. Cabezuelo, C. Abad, A. Perera, J. Prat.

Centre Hospitalari i Cardiològic, Manresa, Barcelona: M. Domènech, J. Badal, J. Malet.

Hospital Universitario de Canarias, La Laguna, Tenerife: R. García-Closas, J. Rodríguez de Vera, A.I. Martín.

Hospital Universitario Nuestra Señora de la Candelaria, Tenerife: J. Taño✝, F. Cáceres.

Hospital General Universitario de Elche, Universidad Miguel Hernández, Elche, Alicante: A. Carrato, F. García-López, M. Ull, A. Teruel, E. Andrada, A. Bustos, A. Castillejo, J.L. Soto.

Universidad de Oviedo, Oviedo, Asturias: A. Tardón.

Hospital San Agustín, Avilés, Asturias: J.L. Guate, J.M. Lanzas, J. Velasco.

Hospital Central Covadonga, Oviedo, Asturias: J.M. Fernández, J.J. Rodríguez, A. Herrero.

Hospital Central General, Oviedo, Asturias: R. Abascal, C. Manzano, T. Miralles.

Hospital de Cabueñes, Gijón, Asturias: M. Rivas, M. Arguelles.

Hospital de Jove, Gijón, Asturias: M. Díaz, J. Sánchez, O. González.

Hospital de Cruz Roja, Gijón, Asturias: A. Mateos, V. Frade.

Hospital Alvarez-Buylla, Mieres, Asturias: P. Muntañola, C. Pravia.

Hospital Jarrio, Coaña, Asturias: A.M. Huescar, F. Huergo.

Hospital Carmen y Severo Ochoa, Cangas, Asturias: J. Mosquera.

### Supplementary Methods and Results

#### Post-GWAS: Functional *in silico* analyses and results

We performed an in-depth, systematic, *in silico* functional analysis at SNP, gene, and pathway levels (**Figure S3**) for variants with a GWAS *p*-value$<1\times10^{-4}$ (N=143). The functional impact of variants and their annotation was performed using DoriTool, an integrative pipeline developed at CNIO, built to combine different bioinformatics algorithms and public databases (Martín-Antoniano, Alonso, Madrid, López De Maturana, & Malats, 2017). We also investigated if the prioritized variants were identified as PC-associated lncRNAs in a catalogue computed using a systems and experimental biology approaches (Arnes et al., 2019).

**eQTL** analysis was done using three independent pancreatic datasets: (1) GTEx version 7 dataset for 220 normal pancreatic tissue samples was obtained from the GTEx Portal on 06/11/2018 (Ardlie et al., 2015); 2) Laboratory of Translational Genomics (LTG) dataset with 95 histologically normal pancreatic samples (Zhang et al., 2018); and 3) The Cancer Genome Atlas (TCGA) dataset with 178 pancreatic tumors (PAAD samples), using the PancanQTL webtool (Gong et al., 2018). RefSeq genes located within +/−500 kb of the marker SNP for each GWAS significant locus were assessed for *cis*-eQTL effects. The eQTL analysis was performed separately for each dataset.

The **mQTL** analysis was based on the methylation and genotyping data from leukocytes DNA generated in 265 controls from the PanGenEU study. Leukocyte DNA methylation data were obtained using the Illumina Infinium MethylationEPIC BeadChip (Illumina, San Diego, CA, USA, 850k) according to the manufacturer's protocol (Infinium HD Methylation Assay). Preprocessing of methylation data included background correction, normalization procedure using the *preprocessQuantile* function of *minfi* (Aryee et al., 2014), and exclusion of 'failed' probes and cross-reactive probes (Pidsley et al., 2016). A linear model was assumed to test the association between each prioritized SNP and the M-value of each individual CpG site 5mC levels. The models were adjusted for age and sex. All analyses were adjusted for multiple testing correction using Benjamini-Hochberg's method (Benjamini & Hochberg, 1995).

For the **pQTL** analysis, we interrogated the genetic atlas of the human plasma proteome including 1,927 genotype-protein associations, recently published (Sun et al., 2018).

**Histone marks** annotation was performed using the DNase hypersensitive sites, histone modifications, transcription start sites (TSS), active promoters, and transcription factor-binding sites in human cell lines and tissues from assays conducted in pancreas tissue catalogued in the ENCODE project (Sloan et al., 2016).

SNPs were mapped to significant **3D chromatin interaction regions** revealed by HOMER (Heinz et al., 2010) using the Hi-C map of pancreas tissue (Schmitt et al., 2016). Then, we annotated those SNPs in the CI region 1 overlapping with an enhancer region, as well as those interacting with a CI region overlapping with a promoter region. Both enhancer and promoter regions were obtained from Roadmap Epigenomics Projects for pancreas (E098) (https://egg2.wustl.edu/roadmap/web_portal/DNase_reg.html#delieation) and were predicted using DNase peaks and core 15-state chromatin state model. Enhancers were linked to genes using the predictions of enhancer-gene links for the Roadmap Epigenomics reference epigenome for pancreas (E098) for states 6, 7 and 12 (http://www.biolchem.ucla.edu/labs/ernst/roadmaplinking/).

We annotated prioritized SNPs in the **differentially opening regions (DORs)** in human pancreatic endocrine and exocrine lineages reported by Arda et al (Arda et al., 2018).

We interrogated one of the largest publicly available collections of genes and variants associated to human diseases using DisGeNET (Piñero et al., 2017), to investigate whether the prioritized variants are also associated with **PC comorbidities or other types of cancer**.

We performed **enrichment analysis at the gene level** to interpret the potential relevance of our prioritized variants in a broader context of genes and molecular pathways. To this end, we used FUMA GWAS platform (Watanabe, Taskesen, Van Bochoven, & Posthuma, 2017),. Moreover, we used DisGeNet R package to analyse the properties of disease genes.

cBioPortal for Cancer Genomics was used to investigate **molecular profiling** at gene level in PC samples and to identify the molecular profiles including mutations and copy number alterations (amplifications/deletions) that our SNP-associated gene set present in PC samples from the four available studies.

Finally, the differential gene **expression profiling** in cancer and normal tissue was evaluated by comparing tumor to normal tissue to find tumor-specific genes using GEPIA, a web-based tool using TCGA, and GTEx data (Ardlie et al., 2015).

We present here the *in silico* functional evidences for the prioritized variants (**Table S3**):

Two variants at **1q21.3** (rs17661062 and rs59942146) in high LD ($r^2$=0.92) and intronic to ***SETDB1*** and ***FAM63A*** were associated with an increased methylation of the cg17724175 in ***MCL1***. *SETDB1* appears altered in 4% of the PC tumors analyzed in cBioPortal.

rs12756006 at **1p22.3** is a regulatory variant in a promoter flanking region which overlaps with a DOR in ductal cells. Moreover, it overlaps with H3K4me1 and H3K9me3 in pancreas.

*NR5A2*-rs4465241 and *NR5A2*-rs3790840 (**1q32.1**) overlap with one and 10 histone marks, respectively (H3K27ac in pancreas; and DNase, H3K27ac, H3K27me3, H3K36me3, H3K4me1, H3K4me3 and H3K9me3 in pancreas and H3K27ac, H3K27me3 and H3K36me3 in endocrine pancreas). As discussed before, variants tagging ***NR5A2*** have previously been associated with PC risk in GWAS. This orphan nuclear receptor participates in a wide variety of processes such as cholesterol and glucose metabolism in the liver, resolution of endoplasmic reticulum stress, intestinal glucocorticoid production, pancreatic development and acinar differentiation, and inflammatory response. In addition, *NR5A2* gene has been previously associated with uric acid levels, also associated with PC and lung adenocarcinoma risk (Cobos et al 2018). Moreover, the expression of *NR5A2* in tumor is significantly decreased compared with its counterpart in normal tissue (Log2=-1.84, adjusted-*p*-value= $1.85x10^{-43}$).

rs6697813 (**1p36.21**) is upstream to ***DDI2*** (*DNA damage inducible 1 homolog 2*) and was significantly associated with differential methylation in leukocytes of four CpG sites annotated in ***CHCHD2P6-RP4-680D5.2***, ***CASP9***, and ***AGMAT-DNAJC16***. The *G* allele is associated with an increased expression of ***C1orf144*** in normal pancreas. Moreover, it overlaps with 7 chromatin marks: two in endocrine pancreatic tissue and five in pancreas tissue. Another *DDI2* variant in linkage equilibrium with our hit has been previously associated with alcoholic chronic pancreatitis (Rosendahl et al., 2018).

*DHRS3*-rs12136952 (**1p36.22**) is an intronic variant of ***DHRS3*** located in an enhancer region of the gene, overlapping a H3K4me1 chromatin mark in pancreas tissue, enriched at active enhancers. This gene is a highly conserved member of the short-chain dehydrogenases/reductases family, which encodes the DHRS3 protein, an endoplasmic reticulum protein (Deisenroth, Itahana, Tollini, Jin, & Zhang, 2011). The variant also overlaps a CTCF binding site and is in a chromatin region contacting the promoter of the *Vacuolar Protein Sorting 13 Homolog D* (***VPS13D***), involved in vesicle transport. This variant is a potential regulator of the expression of *DHRS3*. *DHRS3* appears to be overexpressed in PC tumors versus normal tissue.

The rs11118832 variant at **1q41** is located in an intron of ***DUSP10***. It overlaps with 5 and 6 chromatin marks, respectively. Variants in *DUSP10* were previously reported as associated with colorectal cancer. *DUSP10* is mutated in 1.9% of pancreatic adenocarcinoma/cancer samples in cBioPortal and is differentially expressed in tumor vs normal tissue (Log2= 1.135; *p*-value=7.23E-37).

*SCTR*-rs4383344 (**2q14.2**) is associated with decreased methylation of cg24309134, located in an openSea region. ***SCTR*** codes for the secretin receptor, crucially involved in the function of healthy pancreatic ductal epithelial cells where it stimulates pancreatic bicarbonate, electrolyte, and fluid secretion; its silencing may contribute to tumor growth and progression of PC (Ding, Cheng, McElhiney, Kuntz, & Miller, 2002). Secretin receptors are overexpressed in non-neoplastic pancreas ducts and its isoforms may be correlated with decreased secretin binding in pancreatic ductal tumors.

*KIAA1257*-rs1683813 (**3q21.3**) is associated with a decreased methylation of cg27310733 and with the overexpression of ***LOC653712*** and ***AC112484*.3** in both pancreatic tissue and whole blood (GTEx). Moreover, it overlaps with H3K27me3 in pancreatic tissue. *KIAA1257*-rs9810890 ($r^2$=0.11 with our hit) is associated with dental caries. ***KIAA1257*** is mutated in 0.8% of pancreatic adenocarcinoma/cancer samples in cBioPortal.

rs2584051 (**3p26.2**) is a non-coding transcript exon variant at the *Leucine Rich Repeat Neuronal 1* (***LRRN1***) and intronic of sulfatase modifying factor 1 (***SUMF1***). This regulatory variant overlaps with a DOR in ductal cells and with two chromatin marks associated with active transcription (H3K4me1 and H3K4me3).

*SEC63*-rs12206846 (**6q21**) is associated with an over-methylation of cg00234027. ***SEC63*** encodes a protein involved in protein translocation in the endoplasmic reticulum. The SNP overlaps with H3K36me3 and H3K9me3 in both endocrine and pancreas tissue. An independent variant in this gene (rs11153123) also overlaps with five chromatin marks in pancreas: H3K27ac, H3K36me3, H3K4me1, H3K4me3 and H3K9me3.

*GMDS-AS1*-rs761098 (**6p25.2**) is a regulatory region variant in *GDP-Manose 4, 6-dehydratase Antisense* RNA 1 (Head To Head), a non-coding RNA gene that has been associated with pancreatic ductal adenocarcinoma (Arnes et al., 2019), suggesting that genetic variants therein may contribute to transcriptional regulation in pancreatic cancer. ***GMDS-AS1*** catalyzes the conversion of GDP-mannose to GDP-4-keto-6-deoxymannose, the first step in the synthesis of GDP-fucose from GDP-mannose (using NADP+ as a cofactor). GDP-fucose is the fucose donor used to synthesize all mammalian fucosylated glycans, including ABO blood group antigens, present on the surface of all cell membranes. *GMDS-AS1* appears with a deep deletion in 0.8% of the pancreatic adenocarcinoma/cancer samples in cBioPortal. *GMDS* gene is expressed in PANC-1 cells and in well-differentiated tumors. *GMDS* is enriched in the epithelium in primary tumors ($p$-adj<0.05). A loss-of-function variant in this gene associated with tumor progression. *GMDS* is mutated in two PC samples in cBioPortal and is differentially expressed in PC (TCGA) vs normal tissue (GTEx) (Log2FC=1.625, $p$-value=$1.65 \times 10^{-45}$).

*TNS3*-rs2271311 (**7p12.3**) is associated with an increased methylation of *TNS3*-cg06114556 and overlaps with 5 chromatin marks in pancreas (DNase; H3K27ac; H3K36me3; H3K4me1; and H3K4me3) and 2 in endocrine pancreatic tissue (H3K9ac H3K27ac). The SNP is in linkage equilibrium with rs78417682, a pancreatic cancer susceptibility variant reported in (Klein et al., 2018). ***TNS3*** is altered in 1% of the pancreatic adenocarcinoma cases in cBioPortal.

Two variants at **7q22.1** (rs62484781, and rs62482372; $r^2_{rs62484781\text{-}rs62482372}$=0.7) are located upstream to ***DPY19L2P2***. rs6955512 is associated with increased methylation of SMURF1-cg27297376 and it overlaps with H3K27ac and H3K4me1 marks in both endocrine and pancreatic tissue. Variants in *DPY19L2P2* were previously associated with breast cancer (Michailidou et al., 2017). *DPY19L2P2*-rs62484781 overlaps with H3K27ac and H3K4me1 histone marks in both endocrine and pancreatic tissue. Approximately 2% of PC in cBioPortal showed alterations in ***SMURF1***.

*FBRSL1*-rs6560884 (**12q24.33**) is associated with decreased expression of ***FBRSL1*** in normal pancreatic tissue (GTEx and LTG datasets) and it overlaps with three chromatin marks in pancreas tissue and with one in endocrine pancreatic tissue. This variant is in moderate LD with rs10870474 ($r^2$=0.43), also in *FBRSL1*, which was associated with increased methylation of 5 cpgs (cg25130710; *FBRSL1*-cg03621470, promoter associated; cg15310701, cg15785681 and cg24135151), with an increased expression of the same gene in whole blood (GTEx), and overlaps with H3K27ac, H3K36me3 and H3K9me3 marks in pancreatic tissue. *FBRSL1* is altered in 1.3% of PC tumors.

*PRKCA*-rs11654719 at **17q24.2** is associated with an increased methylation of cg08055746 (OpenSea) and it overlaps with H3K4me1 in both pancreatic and endocrine tissues, and with H3K27ac, H3K4me1 and H3K9me3 marks in pancreatic tissue. ***PRKCA*** participates in the SCTR pathway. Moreover, it is altered in 1.3% of the pancreatic tumors.

rs8111858 (**19p13.3**) is an intronic variant of ***STAP2*** *(Signal-transducing adaptor family member-2*) and *FSD1 (Fibronectin type III and SPRY domain containing 1*). *STAP2* encodes an adaptor protein that regulates various intracellular signaling pathways and promotes tumorigenesis in melanoma and breast cancer (Kitai et al., 2017). This variant seems to be functional because the T allele is associated with lower methylation in leukocytes and with an increased expression of *STAP2* in normal pancreatic tissue. Moreover, it overlaps with three histone chromatin marks which are associated with activation of gene transcription (H3K36me3, H3K4me1 and H3K4me3). *STAP2* is overexpressed in pancreatic tumors in comparison with normal pancreatic tissue.

rs62209634 (**20q11.22**) is an intergenic variant interacting with a chromatin region harboring the promoters of three genes: ***PXMP4***, ***ZNF341,*** and ***RP4-553F4.2***. *PXMP4* is overexpressed in tumors, compared to normal tissue. The SNP is associated with differential expression of ***BPIFB2, CHMP4B*** and ***DYNLRB1*** in normal pancreatic tissue.

No overlapping was found between our 143 variants and pQTLs obtained in the plasma proteome.

No significant enrichment of our gene set was observed for gene expression in normal pancreatic tissue (GTEx v7).

**Shared variants between mental diseases and PC**

PC is one of the tumor entities with one of the highest incidences of depression preceding cancer diagnosis (Eguia, Gonda, & Saif, 2012). To explore the relationship between mental disorders and PC risk, we analyzed the association between variants in genes associated with PC risk in our study and both neuroticism (*RBFOX1* and *PRKCA*) and major depression disorder (*CACNA2D1*, *NRG1*, *NRG1-IT2* and *KB-1047C11.2*) in GWAS catalog and the risk of mental disorders in PC cases from PanGenEU study. None of the associations remained significant after multiple testing correction, the minimum *p*-value being for *RBFOX1*-rs10852686 ($4.95 \times 10^{-03}$).

**Drug repurposing on newly identified genes**

We analyzed whether some of the genes we retrieved either through GWAS-LMI (n=338) or 3D interactions (n=37) were drug-targets. We searched for all these genes detected through LMI and 3D in PharmaGKB database (*downloaded full data on May 28th, 2019* from pharmgkb.org) and searched for their occurrences in that database. We did not find direct evidence of these genes being target for PC current treatments. A total of 23/338 (6.8%) genes in the selected GWAS-LMI distribution were annotated in the list of clinically actionable gene-drug associations for other cancer types or conditions associated with PC. Four genes were already pharmacological targets for several types of cancer (*BCL2*; ovarian neoplasms, *CDKN2A,* and *FOLH1*; lymphoblastic leukemia, *ZNF423*; breast neoplasms) and two genes (*AQP2* and *NBEA*) were clinically actionable to treat type II diabetes mellitus. Intriguingly, ~30% of the actionable genes had associations with drugs to treat mental conditions (*e.g.*, anxiety, depression or psychotic disorders), which have been associated with PC (Eguia et al., 2012). Furthermore, *ALK, BDNF, DDHD1* and *XBP1*, which is 10.8% of the 37 target genes annotated through Hi-C interactions, had at least an entry in the database for any of the categories. Remarkably, *ALK* is already a target for many cancer treatments (neuroblastoma, non-small cell lung cancer and anaplastic lymphoma), and is currently undergoing clinical trial as a target for Ceritinib for treating pancreatic cancer metastasis: clinicaltrials.gov/ct2/show/NCT02227940. *BDNF* is the target for many psychiatric disorders (depression, schizophrenia, substance abuse disorder, among others). *DDHD1* has been associated with warfarin dose-response, and, finally, SNP variants in *XBP1* affect response to platinum compounds in lung cancer treatment.

### REFERENCES

Arda, H. E., Tsai, J., Rosli, Y. R., Giresi, P., Bottino, R., Greenleaf, W. J., … Kim, S. K. (2018). A Chromatin Basis for Cell Lineage and Disease Risk in the Human Pancreas. *Cell Systems*, *7*(3), 310-322.e4. https://doi.org/10.1016/j.cels.2018.07.007

Ardlie, K. G., DeLuca, D. S., Segrè, A. V., Sullivan, T. J., Young, T. R., Gelfand, E. T., … Lockhart. (2015). The Genotype-Tissue Expression (GTEx) pilot analysis: Multitissue gene regulation in humans. *Science*, *348*(6235), 648–660. https://doi.org/10.1126/science.1262110

Arnes, L., Liu, Z., Wang, J., Maurer, H. C., Sagalovskiy, I., Sanchez-Martin, M., … Rabadan, R. (2019). Comprehensive characterisation of compartment-specific long non-coding RNAs associated with pancreatic ductal adenocarcinoma. *Gut*, *68*(3), 499–511. https://doi.org/10.1136/gutjnl-2017-314353

Benjamini, Y., & Hochberg, Y. (1995). Controlling the False Discovery Rate: A Practical and Powerful Approach to Multiple Testing. *Journal of the Royal Statistical Society: Series B (Methodological)*, *57*(1), 289–300. https://doi.org/10.1111/j.2517-6161.1995.tb02031.x

Deisenroth, C., Itahana, Y., Tollini, L., Jin, A., & Zhang, Y. (2011). p53-inducible DHRS3 is an endoplasmic reticulum protein associated with lipid droplet accumulation. *Journal of Biological Chemistry*, *286*(32), 28343–28356. https://doi.org/10.1074/jbc.M111.254227

Ding, W. Q., Cheng, Z. J., McElhiney, J., Kuntz, S. M., & Miller, L. J. (2002). Silencing of secretin receptor function by dimerization with a misspliced variant secretin receptor in ductal pancreatic adenocarcinoma. *Cancer Research*, *62*(18), 5223–5229.

Eguia, V., Gonda, T. A., & Saif, M. W. (2012). Early detection of pancreatic cancer. *JOP : Journal of the Pancreas*, Vol. 13, pp. 131–134. https://doi.org/10.1097/mpa.0000000000001024

Gong, J., Mei, S., Liu, C., Xiang, Y., Ye, Y., Zhang, Z., … Han, L. (2018). PancanQTL: Systematic identification of cis -eQTLs and trans -eQTLs in 33 cancer types. *Nucleic Acids Research*, *46*(D1), D971–D976. https://doi.org/10.1093/nar/gkx861

Heinz, S., Benner, C., Spann, N., Bertolino, E., Lin, Y. C., Laslo, P., … Glass, C. K. (2010). Simple Combinations of Lineage-Determining Transcription Factors Prime cis-Regulatory Elements Required for Macrophage and B Cell Identities. *Molecular Cell*, *38*(4), 576–589. https://doi.org/10.1016/j.molcel.2010.05.004

Kitai, Y., Iwakami, M., Saitoh, K., Togi, S., Isayama, S., Sekine, Y., … Matsuda, T. (2017). STAP-2 protein promotes prostate cancer growth by enhancing epidermal growth factor receptor stabilization. *Journal of Biological Chemistry*, *292*(47), 19392–19399. https://doi.org/10.1074/jbc.M117.802884

Klein, A. P., Wolpin, B. M., Risch, H. A., Stolzenberg-Solomon, R. Z., Mocci, E., Zhang, M., … Amundadottir, L. T. (2018). Genome-wide meta-analysis identifies five new susceptibility loci for pancreatic cancer. *Nature Communications*, *9*(1). https://doi.org/10.1038/s41467-018-02942-5

Martín-Antoniano, I., Alonso, L., Madrid, M., López De Maturana, E., & Malats, N. (2017). DoriTool: A Bioinformatics Integrative Tool for Post-Association Functional Annotation. *Public Health Genomics*, *20*(2), 126–135. https://doi.org/10.1159/000477561

Michailidou, K., Lindström, S., Dennis, J., Beesley, J., Hui, S., Kar, S., … Easton, D. F. (2017). Association analysis identifies 65 new breast cancer risk loci. *Nature*, *551*(7678), 92–94. https://doi.org/10.1038/nature24284

Piñero, J., Bravo, Á., Queralt-Rosinach, N., Gutiérrez-Sacristán, A., Deu-Pons, J., Centeno, E., … Furlong, L. I. (2017). DisGeNET: A comprehensive platform integrating information on human disease-associated genes and variants. *Nucleic Acids Research*, *45*(D1), D833–D839. https://doi.org/10.1093/nar/gkw943

Rosendahl, J., Kirsten, H., Hegyi, E., Kovacs, P., Weiss, F. U., Laumen, H., … Sahin-Tóth, M. (2018). Genome-wide association study identifies inversion in the CTRB1-CTRB2 locus to modify risk for alcoholic and non-alcoholic chronic pancreatitis. *Gut*, *67*(10), 1855–1863. https://doi.org/10.1136/gutjnl-2017-314454

Schmitt, A. D., Hu, M., Jung, I., Xu, Z., Qiu, Y., Tan, C. L., … Ren, B. (2016). A Compendium of Chromatin Contact Maps Reveals Spatially Active Regions in the Human Genome. *Cell Reports*, *17*(8), 2042–2059. https://doi.org/10.1016/j.celrep.2016.10.061

Sloan, C. A., Chan, E. T., Davidson, J. M., Malladi, V. S., Strattan, J. S., Hitz, B. C., … Cherry, J. M. (2016). ENCODE data at the ENCODE portal. *Nucleic Acids Research*, *44*(D1), D726–D732. https://doi.org/10.1093/nar/gkv1160

Sun, B. B., Maranville, J. C., Peters, J. E., Stacey, D., Staley, J. R., Blackshaw, J., … Butterworth, A. S. (2018). Genomic atlas of the human plasma proteome. *Nature*, *558*(7708), 73–79. https://doi.org/10.1038/s41586-018-0175-2

Watanabe, K., Taskesen, E., Van Bochoven, A., & Posthuma, D. (2017). Functional mapping and annotation of genetic associations with FUMA. *Nature Communications*, *8*(1). https://doi.org/10.1038/s41467-017-01261-5

Zhang, M., Lykke-Andersen, S., Zhu, B., Xiao, W., Hoskins, J. W., Zhang, X., …

Amundadottir, L. (2018). Characterising cis-regulatory variation in the transcriptome of histologically normal and tumour-derived pancreatic tissues. *Gut*. https://doi.org/10.1136/gutjnl-2016-313146

**Table S1.** Replication of the SNPs reported as associated with PC risk in European population and published in GWAS Catalog.

| **Gene** | **SNP** | **Chromosome region** | **OR** | ***p*-value** |
|---|---|---|---|---|
| ***WNT2B*** | rs351365 | 1p13.2 | 1.18 | 3.92E-02 |
| ***LOC105371682-RNU6-716P*** | rs10919791 | 1q32.1 | 0.80 | 9.67E-03 |
| ***NR5A2*** | rs2816938 | 1q32.1 | 1.28 | 6.97E-04 |
| ***NR5A2*** | rs3790844 | 1q32.1 | 0.81 | 9.35E-03 |
| ***ETAA1-LOC107985891*** | rs1486134 | 2p14 | 0.91 | 1.74E-01 |
| ***EDNRA*** | rs6537481 | 4q31.22 | 0.86 | 4.21E-02 |
| ***TERT*** | rs2736098 | 5p15.33 | 0.75 | 7.52E-05 |
| ***CLPTM1L*** | rs401681 | 5p15.33 | 1.18 | 9.47E-03 |
| ***CLPTM1L*** | rs31490 | 5p15.33 | 1.18 | 8.85E-03 |
| ***TSN3*** | rs73328514 | 7p12.3 | 0.77 | 1.05E-02 |
| ***LINC-PINT*** | rs6971499 | 7q32.3 | 0.79 | 2.04E-02 |
| ***HNF4G*** | rs294147 | 8q21.11 | 1.14 | 4.60E-02 |
| ***SMC2*** | rs10991043 | 9q31.1 | 1.14 | 4.50E-02 |
| ***RNY1P8-MARK2P12*** | rs9543325 | 13q22.1 | 0.85 | 9.96E-03 |
| ***BCAR1*** | rs7190458 | 16q23.1 | 1.62 | 4.14E-04 |
| ***LINC00673*** | rs11655237 | 17q24.3 | 1.46 | 1.43E-04 |
| ***LINC00673*** | rs7214041 | 17q24.3 | 1.47 | 9.12E-05 |

**Table S2.** Validated variants (at the nominal p-value) in PanScan and PanC4 populations, among the top 20 SNPs identified in the PanGenEU GWAS study.

| Cytoband | SNP | $I^2$ | OR | 95%CI | *p-value* |
|---|---|---|---|---|---|
| 17q24.3 | *LINC00673*-rs7214041 | 10.70 | 1.24 | (1.19,1.31) | 9.63E-20 |
| 5p15.33 | *TERT*-rs2736098 | 0.00 | 0.82 | (0.77, 0.86) | 1.29E-13 |
| 1q32.1 | *NR5A2*-rs4465241 | 30.07 | 0.81 | (0.76, 0.87) | 3.27E-10 |
| 1q32.1 | *NR5A2*-rs3790840 | 58.07 | 1.23 | (1.12, 1.34) | 5.91E-06 |
| 19p13.11 | *NWD1*-rs773914 | 80.37 | 1.27 | (1.02, 1.58) | 3.39E-02 |
| 10p11.22 | rs793088 | 82.68 | 1.12 | (1.01, 1.25) | 3.70E-02 |
| 12q22 | *NDUFA12*-rs249184 | 85.34 | 1.25 | (1.00, 1.55) | 4.93E-02 |

The following Supplementary Tables are provided in separate files.

**Table S3**. Annotation and functional *in silico* analysis of the 143 prioritized SNPs (.xls file).

**Table S4.** Results from the gene enrichment analysis performed with FUMA (.txt file).

**Table S5.** List of the 624 SNPs prioritized according to their local Moran index (LMI) (.xls file).

**Table S6.** List of the 76 SNPs overlapping with a chromatin interaction region (bait regions) and their 54 targets in the chromatin interaction regions (.xls file).

**Table S7.** Characteristics of the study populations.

| | PanGenEU+EPICURO | | PanScan I+II | | PanScan III | | PanC4 | |
|---|---|---|---|---|---|---|---|---|
| | Cases (N=1317) | Controls (N=1616) | Cases (N=3525) | Controls (N=3642) | Cases (N=1582) | Controls (N=5203) | Cases (N=3933) | Controls (N=3651) |
| Age (mean, SD) | 64.2 (12.4) | 64.5 (11.5) | 66.2 (10.1) | 66.6 (10.2) | * | * | | |
| Sex | | | | | | | | |
| Male | 575 (43.66%) | 391 (24.20%) | 1841 (52.2%) | 1885 (51.8%) | 781 (49.4%) | 3866 (74.3%) | 2280 (58.0%) | 2038 (55.8%) |
| Female | 742 (56.34%) | 1225 (75.80%) | 1684 (47.8%) | 1757 (48.2%) | 801 (50.6%) | 1337 (25.7%) | 1653 (42.0%) | 1613 (44.2%) |
| Region[a] | | | | | | | | |
| 1 | 32 (2.43%) | 47 (2.91%) | | | | | | |
| 2 | 1045 (79.35%) | 1513 (94.63%) | | | | | | |
| 3 | 240 (18.22%) | 56 (3.47%) | | | | | | |

a: EPICURO control population
*Age for PanScan III cases and controls was provided as age groups (10 year). See Wolpin *et.al* Nature Genetics 2014

**Figure S1**. GWAS Manhattan plot for the PanGenEU study.

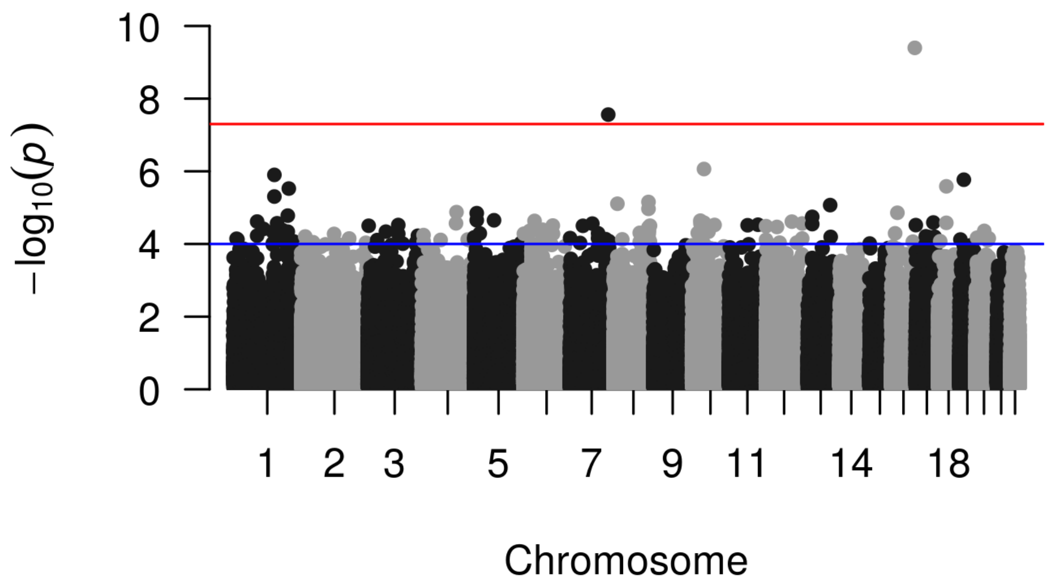

**Figure S2**. Q-Q plots for the PanGenEU (S2a) and PanGenEU&EPICURO study populations (S2b).

**Figure S2a**
(λ=1.04)

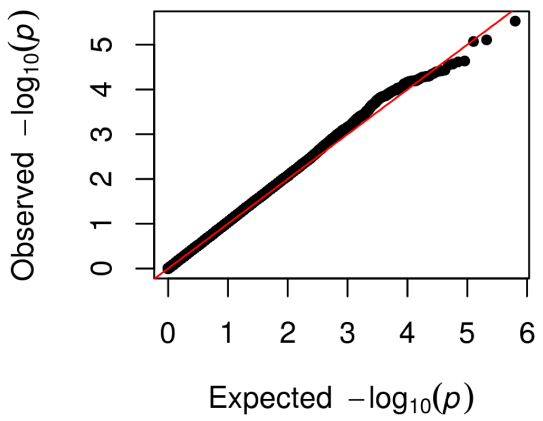


**Figure S2b**
(λ=1.08)

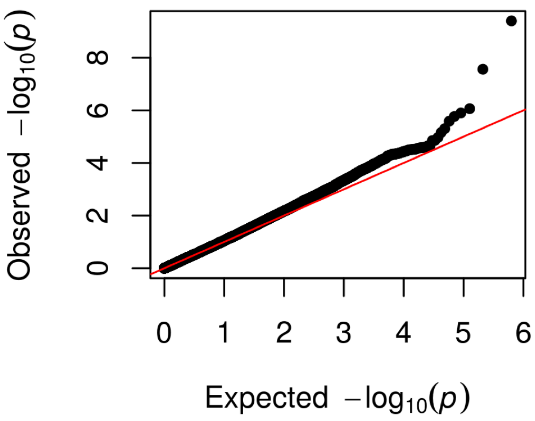

**Figure S3**. Functional *in silico* analysis strategy.

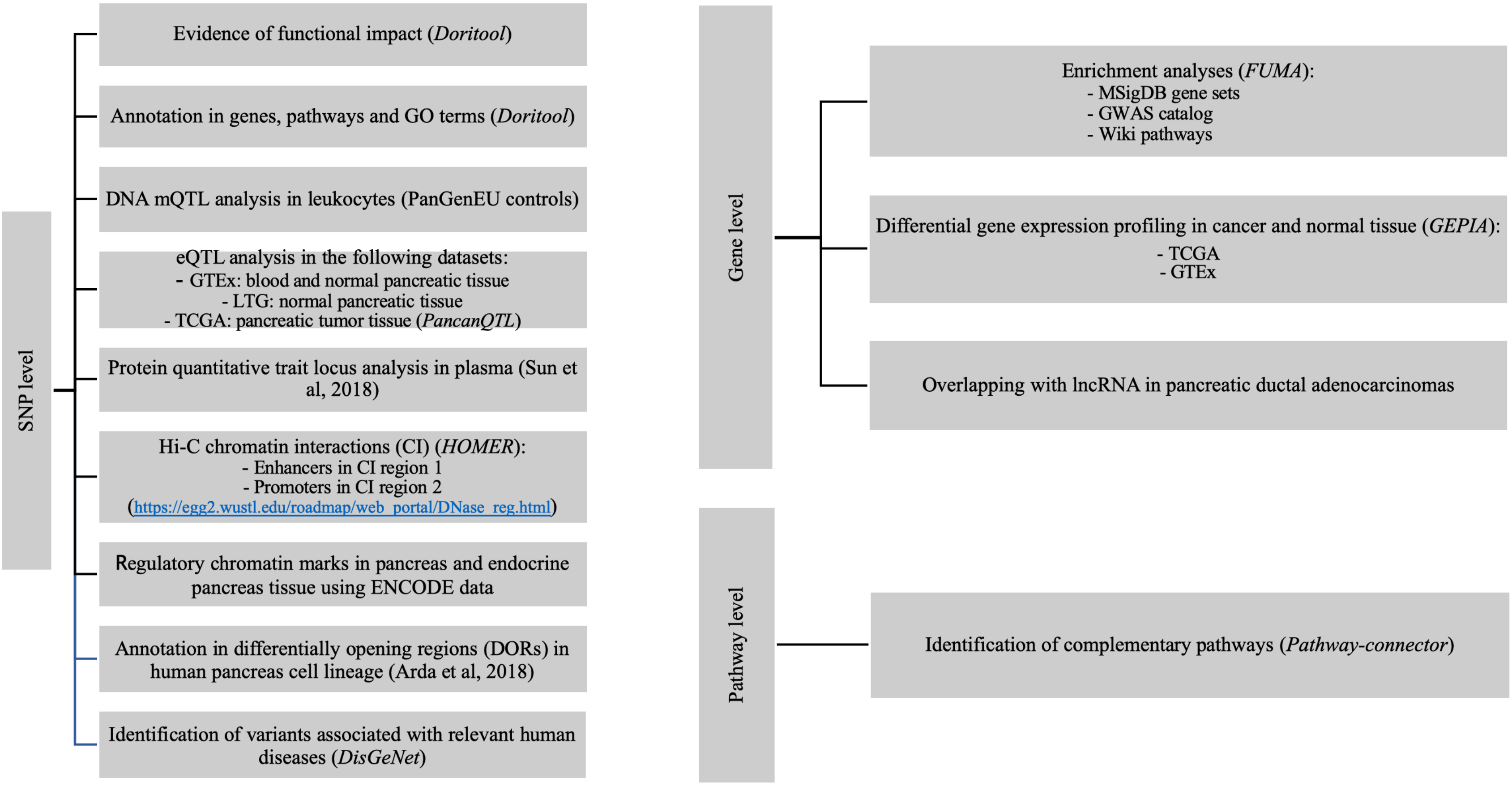

**Figure S4**. Complementary NETWORK (in blue our input pathways; in green, the complementary nodes included) obtained with Pathway-connector webtool.

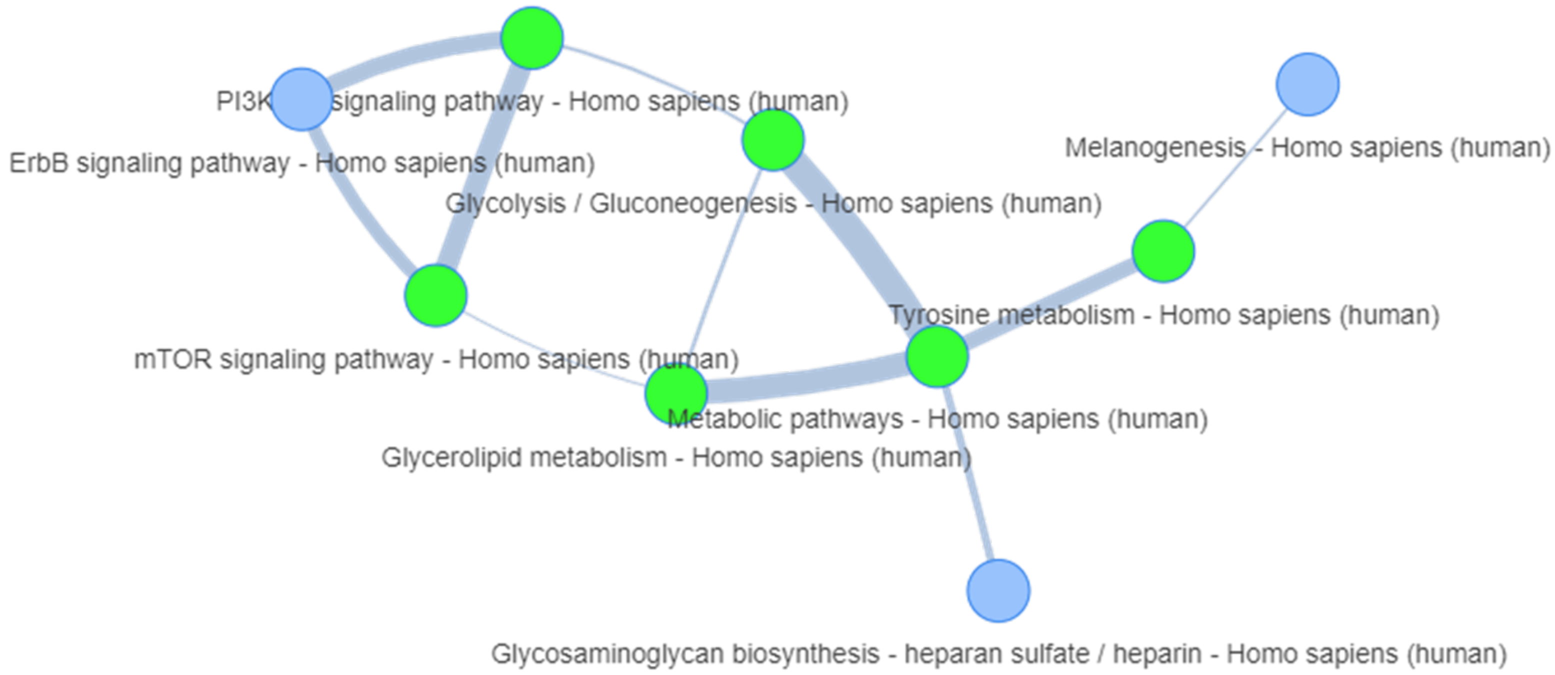

**Figure S5**. Minor allele frequency (MAF) distributions for the top 97 SNPs identified by LMI (in pink) and by GWAS (in blue).

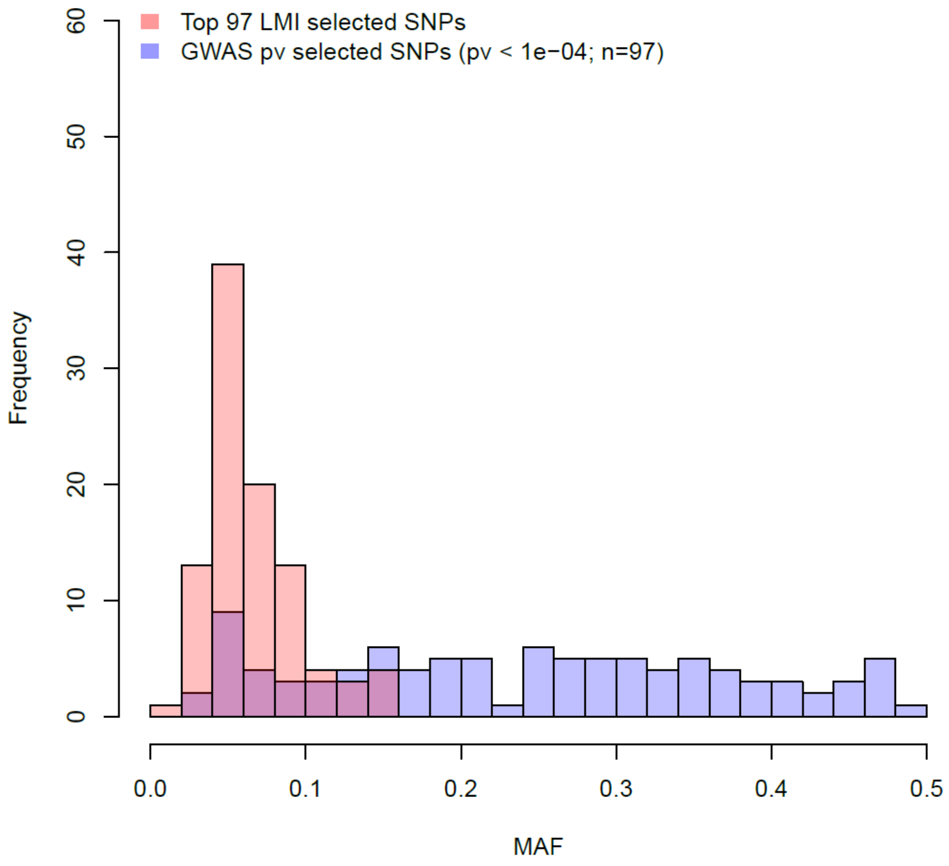

**Figure S6**. Results of the benchmarking test showing that the median rank position of the LMI values for the 22 PC signals from the GWAS Catalog is significantly higher than 10,000 randomly selected sets of the same size.

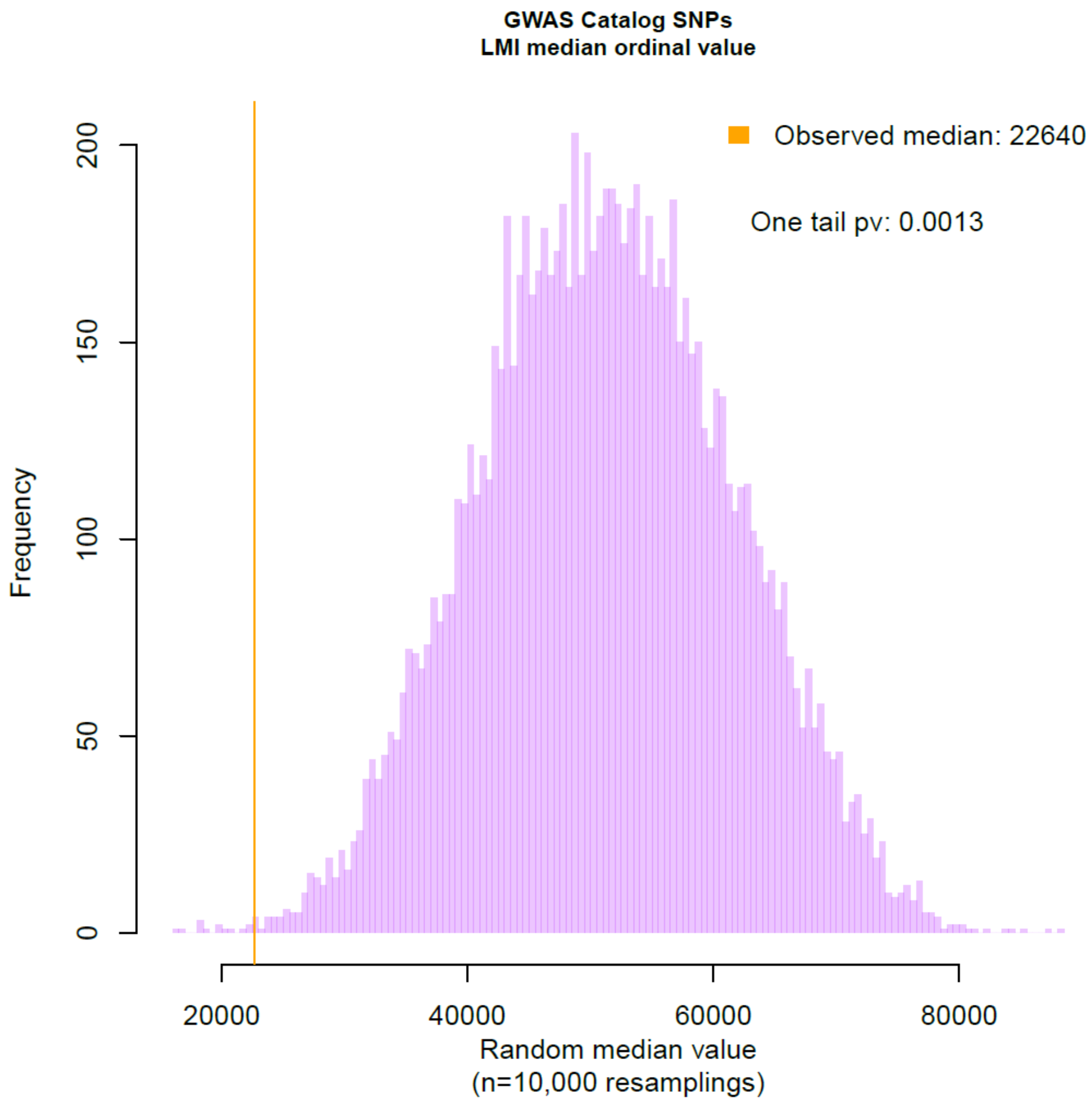
